# Experiment-guided AlphaFold3 resolves accurate protein ensembles

**DOI:** 10.1101/2025.10.11.681796

**Authors:** Advaith Maddipatla, Nadav Sellam Bojan, Meital Bojan, Volodymyr Masalitin, Sanketh Vedula, Paul Schanda, Ailie Marx, Alex M. Bronstein

## Abstract

AlphaFold3 predicts highly accurate protein structures from sequence, but tends to collapse to a single dominant conformation, even when the underlying structure is inherently heterogeneous. Moreover, its predictions are oblivious to experimental conditions that can alter local sequence conformation. In this work we show that AlphaFold3 can be guided to match data obtained by NMR spectroscopy, X-ray crystallography and cryo-EM experiments, and combinations thereof. Our approach can also incorporate data that explicitly report on dynamics, such as site-resolved order parameters. We demonstrate that this methodology can generate ensembles of conformations having less distance restraint violations than traditionally resolved NMR structures and uncover unmodelled alternate conformations detectable in electron density. This methodology paves the way for the development of experimentally aware predictive models that capture the ensemble nature of protein structures.

## Introduction

Proteins are inherently dynamic entities, sampling variable conformational states in response to their surroundings and to fulfill their biological roles. While experimental techniques including X-ray crystallography, NMR spectroscopy and cryo-electron microscopy observe ensemble averages of these macromolecular structures, protein structural models often only report the dominant conformation, overlooking the underlying conformational heterogeneity.

Leveraging on the fundamental discovery that amino acids in spatial proximity within a protein structure coevolve (Göbel et al., 1994; Hopf et al., 2014), and training on the hundreds of thousands of experimentally determined X-ray crystal structures available today, deep learning-based models such as AlphaFold (Jumper et al., 2021; Abramson et al., 2024), have revolutionized structural biology by enabling protein structure prediction approaching experimental accuracy. However, AlphaFold’s training objective – to predict a single “most probable” structure – biases its output toward static snapshots, effectively marginalizing conformational heterogeneity encoded in its training data. This emphasizes the need for new models that can explicitly model protein ensembles that are faithful to experimental measurements.

To bridge this gap, we present a framework for building experiment-grounded protein structure generative models that infer conformational ensembles consistent with measured experimental data. The key idea is to treat state-of-the-art protein structure predictors (e.g., AlphaFold3) as sequence-conditioned structural priors, and cast ensemble modeling as posterior inference of protein structures given experimental measurements.

Recent studies have explored ways to incorporate experimental restraints into structure prediction. AlphaLink (Stahl et al., 2023, 2024) integrates crosslinking mass-spec–derived distances as bias terms in AlphaFold’s pair representation to guide single-structure inference. Here we instead modify the generative process of AlphaFold3 itself: experimental likelihoods enter as guidance terms during sampling, enabling direct conditioning on any experimental modality to generate ensembles consistent with both sequence and experiment.

## Results

We adapt AlphaFold3 to generate ensembles by modifying the reverse diffusion steps such that it includes a gradient-based guidance term derived from the experimental likelihood. This term introduces a data-dependent force that steers sampling toward structures compatible with measurements. A scaling hyperparameter controls the influence of this guidance, allowing interpolation between purely prior-driven and strongly data-conditioned outputs. Our framework supports multiple experimental modalities by defining appropriate differentiable likelihood functions. Following sampling, we perform energy minimization to correct geometric distortions and ensure chemical plausibility – a procedure that is used in many generative models including AlphaFold itself. We then apply ensemble selection to identify the minimal subset of structures that best explain the experimental observations. Refer to Figure 1 for a schematic description of the proposed method and to the Supplementary Methods section in the SI for algorithmic and implementation details.

**Figure 1.**
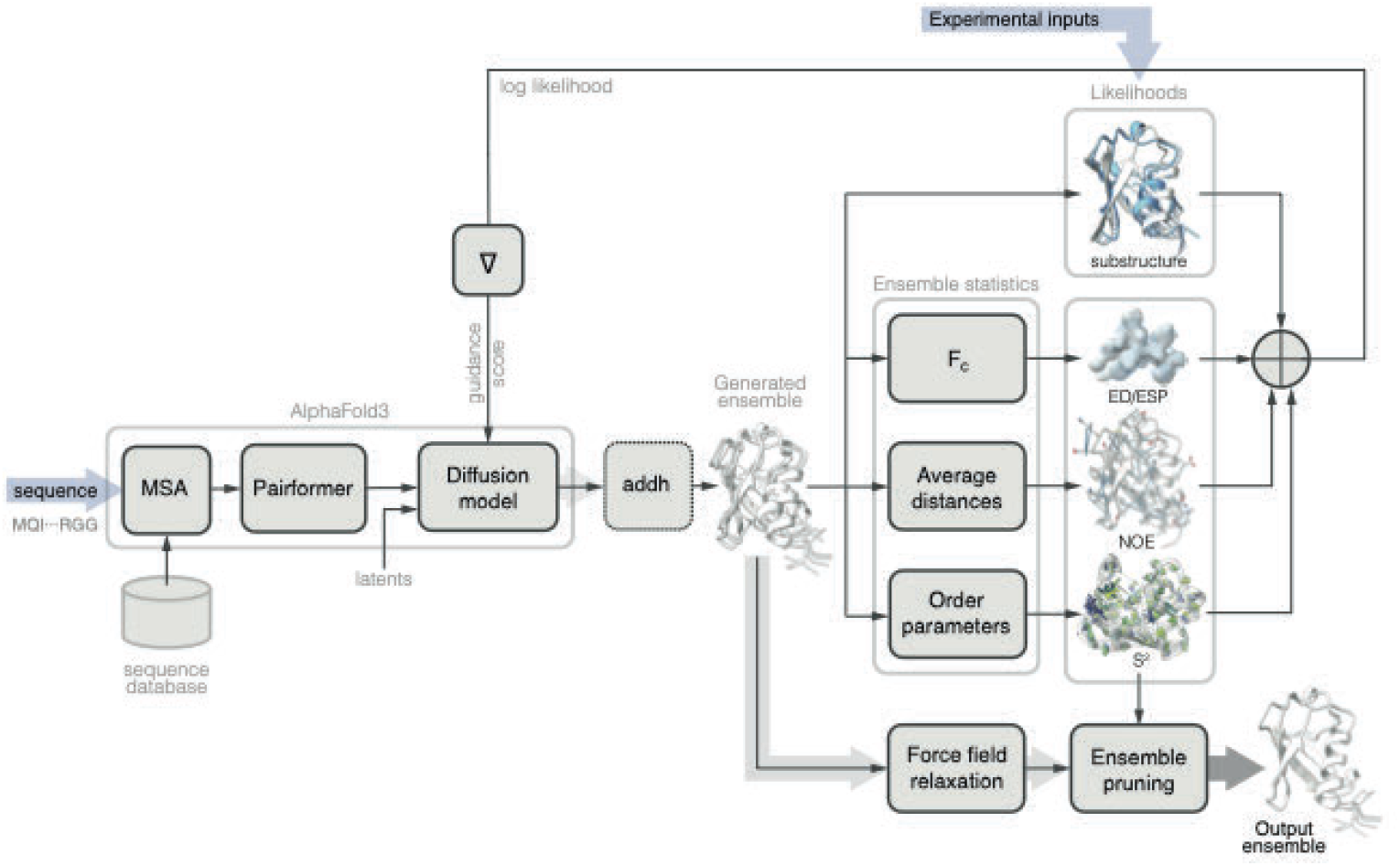
Schematic depiction of the proposed method. AlphaFold3 allows the sampling of protein structures given an amino acid sequence. To further condition the model by experimental observations, at each time step of the AlphaFold3 diffusion model, an ensemble of structures is generated. Likelihoods of experimental observations are calculated given each individual ensemble member (e.g., to enforce locations of certain atoms) and on ensemble averages (e.g., calculated electron density Fc, average inter-atomic distances, or order parameters). The gradient of the combined log-likelihood terms is used as the guidance term. At the final diffusion step, the generated ensemble is refined by force field relaxation and pruned to select a minimal subset maximizing the likelihood terms.

### Guiding AlphaFold3 with NMR distance restraints and order parameters efficiently generates experimentally accurate ensembles

NMR structure determination is largely based on the detection of inter-atomic proximities, which is a manifestation of the dipolar coupling between spins, and detected in Nuclear Overhauser Effect (NOE) experiments in solution-state NMR (Shcherbakov et al., 2021). Typically the structure is then determined by applying these distance restraints in molecular dynamics (MD) simulations to obtain the set of lowest energy conformations (Wang et al., 2004; Schwieters et al., 2006; Güntert, 2004). However, simulating conformers individually often leads to mode collapse, producing rigid ensembles that poorly capture true conformational dynamics. Also, oftentimes no single mode’s pairwise Euclidean distances can fully explain the observed distance averages over multiple incongruent conformations. The explicit determination of multiple state-structural ensembles is becoming more accessible (Vögeli et al., 2016, Chi et al., 2015), but it often requires highly accurate (“exact”) NOE distances, and is not yet common practice in NMR. Ensemble-based MD approaches simulating multiple conformers simultaneously to satisfy experimental restraints remain computationally expensive, requiring days even for small systems like the 76-residue ubiquitin (Lindorff-Larsen et al., 2005 and Lange et al., 2008).

We use ubiquitin (PDB 1D3Z, BMRB 6457, 5387) as an NMR structure determination benchmark. Guiding AlphaFold3 with NOE distance restraints generates ensembles with significantly less pairwise distance constraint violations. Fig. 2A shows the cumulative distribution of violation magnitudes; a spatial visualization of the violated constraints are depicted in Fig. 2C. From Fig. 2B, it is evident that the ensemble produced by NOE-guided AlphaFold3 is more heterogeneous. SI Figs. 1 and 2 evaluate the improvement of constraint satisfaction with the increase of the relative weight of the NOE guidance term. The runtime of the proposed method is several minutes, overcoming the computational bottleneck of traditional ensemble methods.

**Figure 2.**
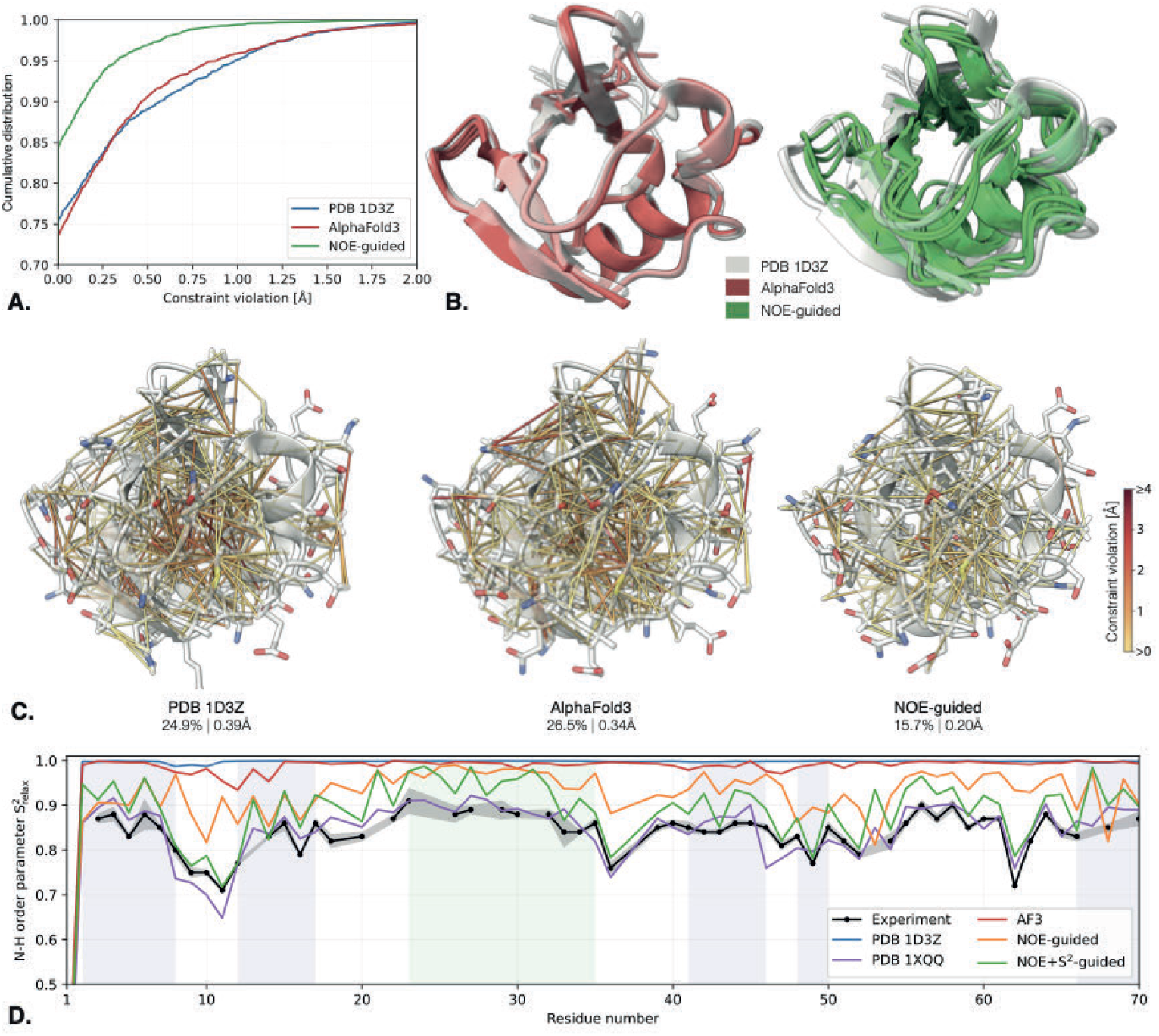
NMR-guided AlphaFold3. Guidance of ubiquitin structure by NOE pairwise distance bounds enforced as ensemble statistics produces significantly less constraint violations than the PDB structure ensemble 1D3Z. (A) Cumulative distribution of constraint violations; (B) unguided (red) and NOE-guided (green) ensembles overlaid on the PDB structure (transparent white); (C) visualization of the pairwise distance constraint violations for the PDB ensemble, unguided and NOE-guided predictions. The fraction of the violated constraints (out of total 1212) and their median violation are reported below each structure. A single best-fitting structure from the ensemble is shown for clarity; (D) Relaxation amide order parameter S^2^ from Lienin et al., 1998 (black) vs. order parameter calculated from the ensembles of different methods. Explicitly guiding the ensemble with NOE and ^15^N-relaxation derived S^2^ constraints (green) produces good agreement with the experiment (r=0.93, q=0.06) and is comparable to the PDB ensemble 1XQQ (purple) fitted to match structure and dynamics (r=0.87, q=0.04). For full results with relaxation and residual dipolar coupling (RDC) data, refer to Supplementary Figures S2, S3.

As an independent verification of ensemble fit to actual protein dynamics, we cross-validated our results using experimental spin relaxation measurements of the backbone amide bond order parameter (S^2^) on time scales shorter than a few nanoseconds (Palmer, 2004). We computed ensemble order parameters and compared them to experimental measurements reported in (Lienin et al., 1998). As illustrated in Fig. 2D, the deposited ensemble derived from standard NMR protocols (PDB 1D3Z) as well as the unguided AlphaFold3 prediction are dominated by rigid conformations. Additional quantitative analysis in SI Fig. 3 shows that these ensembles exhibit only moderate agreement with the experimental S^2^. In contrast, NOE-guided AlphaFold3 significantly improves agreement, better replicating the dynamic behavior observed in both flexible and structured regions.

In order to better match the N-H order parameter experimental data, we added an S^2^ guidance term to the NOE-guided AlphaFold3. This resulted in a significantly higher agreement with the experimental S^2^ measurements and comparable to the result of computationally expensive NMR-guided ensemble MD simulations (PDB: 1XQQ from Lindorff-Larsen et al., 2005) at a small fraction of the computational cost (Fig. 2D and SI Fig. 3A-C). For the case of ubiquitin, a number of additional experimental data have been collected, including methyl axis order parameters from spin relaxation (reporting on ps-ns motions, Liao et al., 2012) and order parameters from residual dipolar couplings for amide and methyl sites, which report on ps-ms motions (Markwick et al., 2009; Sabo et al., 2014; Lange et al., 2008). SI Figs. 3D-F and 4 show the result of guidance with these data. In all cases guidance with NOE+S^2^ terms shows remarkably high agreement with the experiment at no or little expense of pairwise distance constraint violation.

We further evaluated our method on two benchmarks: 8 peptides shown to be mispredicted by AlphaFold3 (Vedula et al. 2025, McDonald et al. 2023) and a subset of 83 proteins from the recently compiled 100 NMR spectra database (Klukowski et al., 2024). SI Table 1 details quantitative performance indicators for individual structures, while Fig. 5A presents summary statistics. We observe that compared to the ensembles deposited into the PDB, NOE-guided AlphaFold3 improves distance constraint satisfaction in 70 out of 91 cases (about 77%), compared to unguided AlphaFold3 which outperforms the PDB ensembles only in 15 cases (17%). NOE-guided AlphaFold3 outperforms its unguided counterpart in all cases. Additional comparison to inductive ensemble generative models including AlphaFlow and ESMFlow (Jing et al., 2024), BioEmu (Lewis et al., 2025) and AFCluster (Wayment-Steele et al., 2024) is presented in SI Table 6.

In all the reported cases, we used uniformly weighed ensembles. SI Figure 11 presents an additional evaluation when a force-field predicted energy is used to weigh the different conformations either as a post-processing step or by fully integrating the force field into the guidance term and producing energy-weighed ensembles maximizing the experimental likelihood. The latter weighing technique improves the distance constraint satisfaction in 78 cases (about 87%) with the median improvement of about 20%. For methodological details, refer to section 2.13 in the Supplementary Methods.

Fig. 3 depicts examples of ensembles generated by NOE-guided AlphaFold3. We observe that AlphaFold3 tends to over-predict order (e.g., longer helical structures in 2B3W, 2JVD and 6SOW). In these cases, NOE guidance increases the ensemble heterogeneity, considerably improving constraint satisfaction and outperforming the corresponding PDB ensembles. In 2K52, no single conformer of the ensemble satisfies the distance constraints, while the ensemble average guidance term significantly improves constraint satisfaction, also compared to the PDB ensemble. In three other cases, AlphaFold3 completely mispredicts parts of the structure (2LI3, 2B3W and 1DEC) resulting in many distance constraint violations. NOE guidance significantly reduces the number of violations.

Finally, as an independent source of validation, we used the ANSURR metrics introduced in (Fowler et al., 2020) that compare the rigidity profile of the model to that inferred from NMR chemical shifts. We observe that ensembles generated by NOE-guided AlphaFold3 improve the RMSD score in 92% of the cases and the correlation score in 54% of the cases when compared to the corresponding PDB ensembles. Refer to SI Fig. 5 for more information.

**Figure 3.**
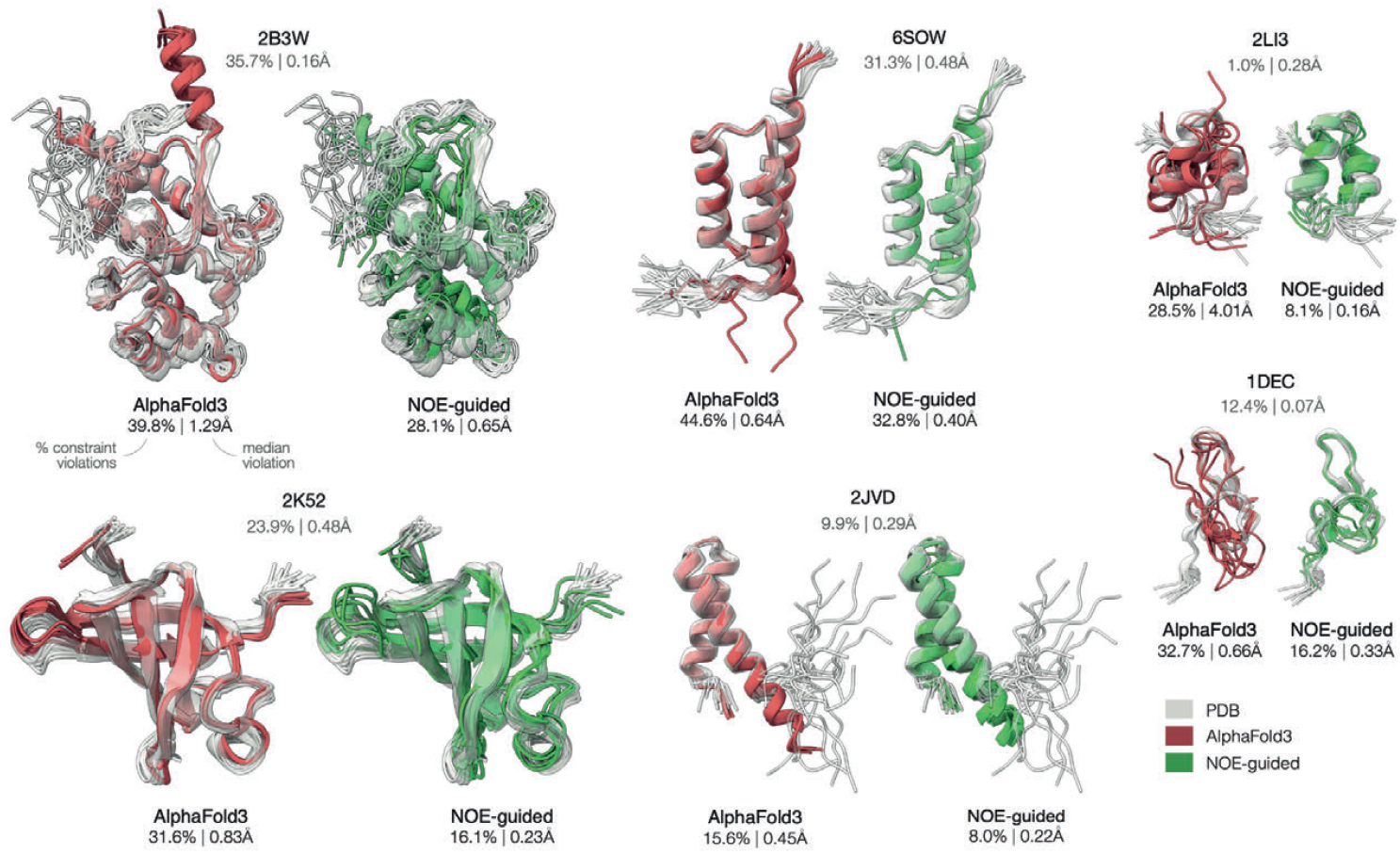
Conformation ensembles generated for six proteins. using unguided AlphaFold3 (red) and its NOE-guided version (green). Ensembles are overlaid on corresponding NMR structures solved from the same NOESY data shown in transparent white. PDB identifiers are indicated above each structure. The numbers below report the percentage of violated constraints and their median violation.

### Guiding AlphaFold3 with electron density captures previously unmodelled residues in X-ray crystallography data

AlphaFold3 is oblivious to environmental influences that may be present in experimental settings, such as interactions with ions, ligands and other macromolecular partners or different crystal contacts resulting in conformational variations. For example, heat shock protein 90-alpha crystallises as a dimer in the asymmetric unit cell with only chain A, and not chain B, captured in the ligand bound state (PDB 6CYH). The loop adjacent to this ligand adopts a significantly different conformation between the two chains, neither of which is accurately predicted by AlphaFold3. Fig. 4A shows that the poor AlphaFold3 prediction is restored to experimental accuracy by electron density guidance. The method is similarly successful when applied across experiments. SI Table 2 shows that a pair of sequence identical constructs for the Orf9b protein from SARS-CoV-2 exhibit varying conformations at several protein chain locations (PDB 9N55 and 9MZB). Electron density guidance successfully restores the AlphaFold3 model generation to match the PDB deposited structures. The success of fitting on electron density maps having very different resolutions (PDB 9N55 at 1.65Å and 9MZB at 2.8Å) pays tribute to the generalizability of our method. In yet another example, we consider a pair of myoglobin structures that differ in surface-exposed loop conformations due to altered crystal packing (PDB 1U7R and 1U7S). Electron density guided AlphaFold reproduces both conformations with density fits comparable to deposited models. Interestingly, it has been noted that these alternate conformations closely resemble those observed in solution by NMR (PDB 1MYF), suggesting that the lattice selects from intrinsically adopted conformations (Kondrashov et al., 2008) and underscores the biological relevance of training AlphaFold on ensembles detectable in crystal density, now an achievable task with our methodology.

**Figure 4.**
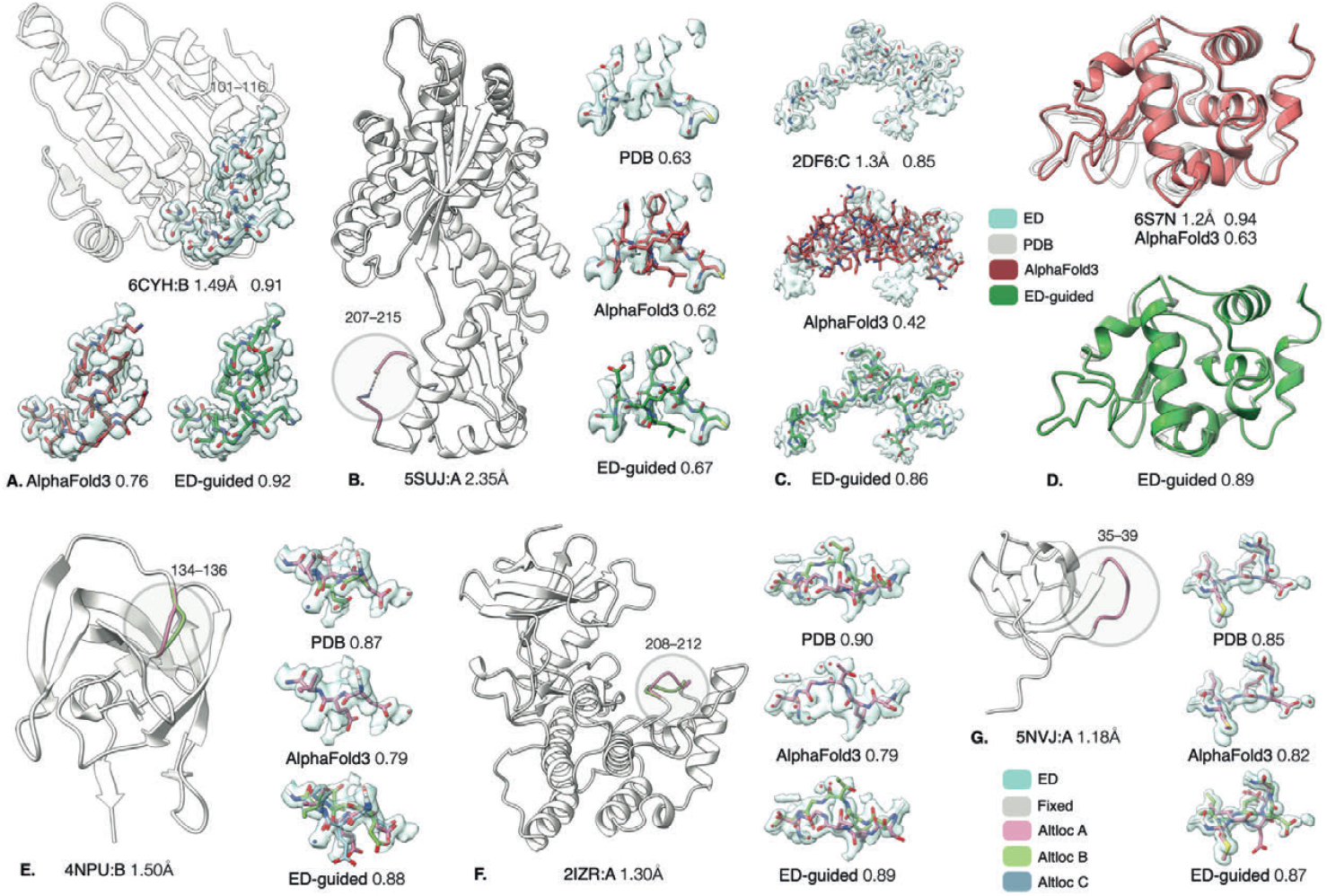
X-ray electron density (ED)-guided Alphafold produces structures and ensembles of equivalent or better accuracy to those deposited in the PDB and detects plausible conformations in unmodelled electron density. (A) 6CYH chain B is poorly predicted by AlphaFold3 in the region of amino acids 101–116. Guidance using the electron density maps restores the fit of the model. (B) An unmodelled region in 5SUJ has different conformations at two locations due to the presence of ligands in one chain. (C) The structure of an 18 amino acid peptide from PAK2 bound to the SH3 Domain of betaPIX is poorly predicted by AlphaFold3 but this prediction can be guided by the electron density to recapture the experimental structure. (D) AlphaFold3 poorly predicts multiple regions of 6S7N. ED guidance essentially reproduces the PDB structure. (E-F) ED-guided AlphaFold3 produces conformational ensembles explaining the multi-modal electron density on par or better than the PDB model. (G) ED-guided AlphaFold3 predicts unmodelled altlocs in 5NVJ better explaining the electron density. Electron densities are visualized as 0.5e^−^/Å^3^-isosurfaces. Numbers below indicate cosine similarities between calculated and observed densities.

**Figure 5.**
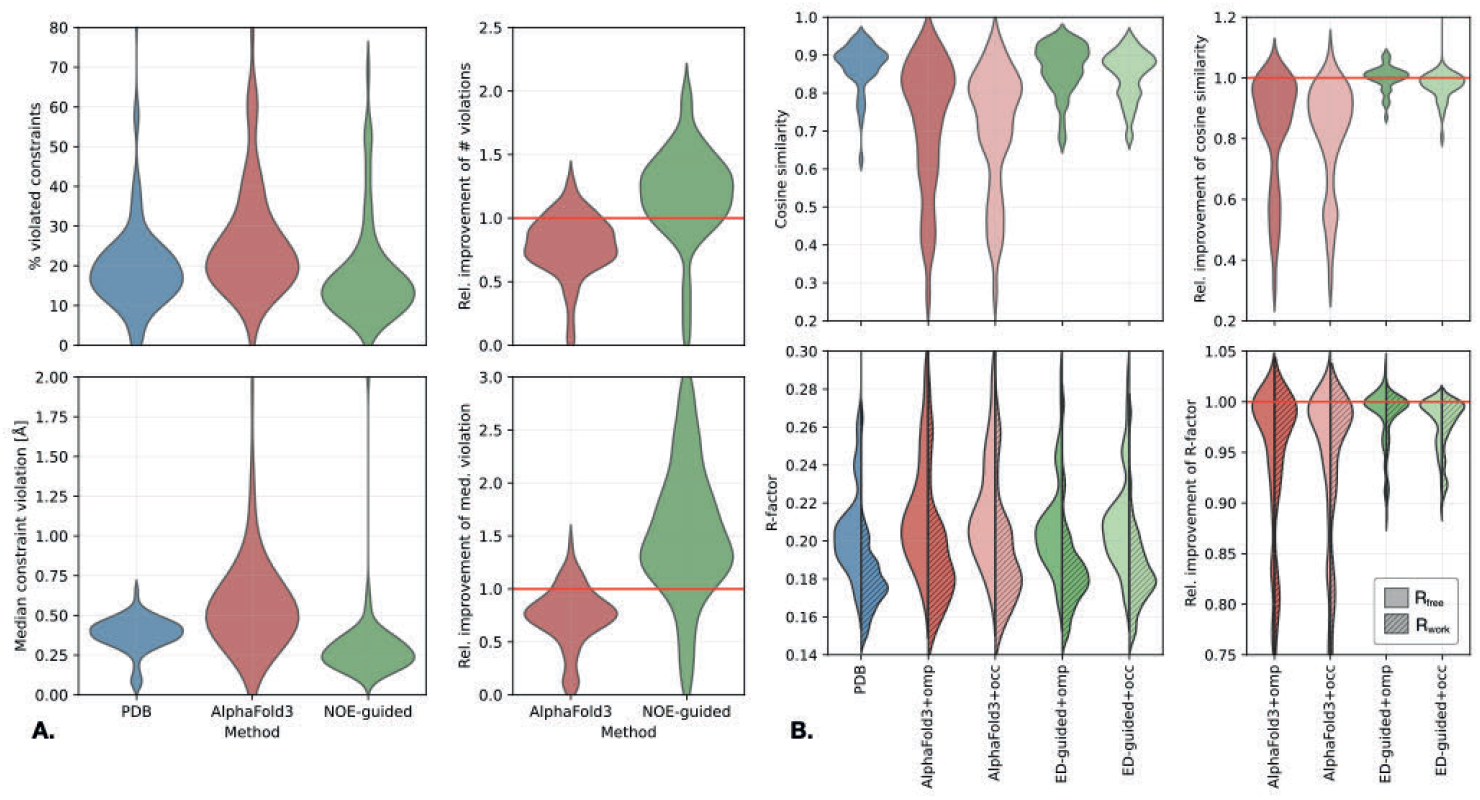
Summary statistics on evaluated NMR and X-ray crystallographic benchmarks. (A) Distribution of % of violated distance constraints (top left) and median violation (bottom left) for PDB, unguided and NOE-guided AlphaFold3 and the relative improvement of these metrics compared to the deposited PDB structures. (B) Distribution of F_o_ and F_c_ cosine similarity (top left) and R-factors (top right, split between R_free_ and R_work_), with the relative improvement shown on the right. Additional summary statistics are shown in SI Figures 10 and 11.

Another region-specific phenomenon in X-ray crystal structures are unmodelled regions. Local flexibility results in a smeared electron density map (measured by the B factor), to the point where the density is too blurred to readily infer structure. We show in Fig. 4B that density guided AlphaFold3 has the potential to fill in missing regions to best explain the scarce density. The crystal structure of an uncharacterized protein LPG2148 from Legionella pneumophila (PDB 5SUJ) crystallizes as a dimer in the asymmetric unit cell and both chains have a missing segment in the model at the same surface exposed loop region. Fig. 4B and SI Fig. 7 show that the density guided version fills the missing space in both chains with better local real space correlation to the sparse density than the AlpaFold3 prediction or the commonly used PDBfixer (Eastman *et al*., 2017).

Another challenge in crystallography is modelling peptides. These short protein chains are often very poorly predicted by AlphaFold3 since they are highly adaptable to their binding partners. Fig. 4C shows that our density-guided method restores the fit of the peptide to the electron density from a very poor AlphaFold3 prediction. This example leverages AlphaFold3 as a simple structural prior, since these ensembles were generated without MSA conditioning. Further examples in SI Table 2 and the summary statistics in Fig. 5B highlights the method’s ability to generalize across sequences of varying lengths. Comparison to ensemble generative models is presented in SI Table 7. Going beyond segment based guidance we automatically guide an entire protein chain through an iterative process of detection and improvement of regions poorly fit to the electron density and phase refinement. The final fitted structure shown in Fig. 4D results from iterative improvements in real space correlation (SI Fig. 8). We hypothesize that the results appear slightly inferior to those deposited in the PDB mainly due to the manual placement of ions and structural waters that AlphaFold does not generate.

Although early crystallographic refinements prioritized single-conformer models, often protein crystal structures can also capture ensembles as two or more distinct conformations within the same electron density map (Smith et al., 1986; Furnham et al., 2006; van den Bedem & Fraser, 2015; Wankowicz et al., 2024). A recent study by Rosenberg et al. (2024a) compiled a comprehensive catalog of altlocs from PDB structures and also showed that even for regions with well-separated and stable altlocs, structural ensemble predictors such as AlphaFold3 fail to reproduce the experimentally determined distributions or capture the bimodal nature of backbone conformations (Rosenberg et al., 2024b). Here, we demonstrate that electron density guidance on this same set of altlocs generates an ensemble that captures the backbone’s bi-modal nature, providing a better explanation of the density. This method is effective for different degrees of backbone separation spanning from 1Å (Fig. 4E) to tens of Ångstroms (Fig. 4F). SI Table 2 further demonstrates that alternate conformations are successfully detected across different sequence types and lengths and within electron density maps having different resolutions and qualities. Moreover, SI Table 4 shows that these conformations are consistently reproduced across independent runs using different types of density maps, with the consistency further improving as batch size increases (SI Fig. 9). Finally, we demonstrate the potential of this method for detecting previously unmodelled altloc conformations (Fig. 4G).

### Guiding AlphaFold3 with cryoEM maps

CryoEM imaging allows to capture the conformations of intricate protein complexes in the form of electrostatic potential (ESP) maps. Since the standard reconstruction pipelines rely on rigid body consensus sets, individual ESP maps do not capture the true thermodynamic heterogeneity of the particle. At the same time, the achievable resolution – especially in flexible regions – is often insufficient for precise placement of individual atoms. We demonstrate on several examples that AlphaFold3 guidance with ESP maps achieves better *ab initio* modeling accuracy.

Fig. 6A shows the reconstruction of insulin receptor IR-B in its symmetric apo state (PDB 8U4B, EMDB 41877) and asymmetric conformation (PDB 8U4E, EMDB 41880) with three bound IGF2 factors. AlphaFold3 tends to over-symmetrize the predicted homodimers, mispredicting large portions of the asymmetric conformation but also inaccurately predicting the symmetric parts. ESP map guidance produces conformations that agree with the observed densities.

**Figure 6.**
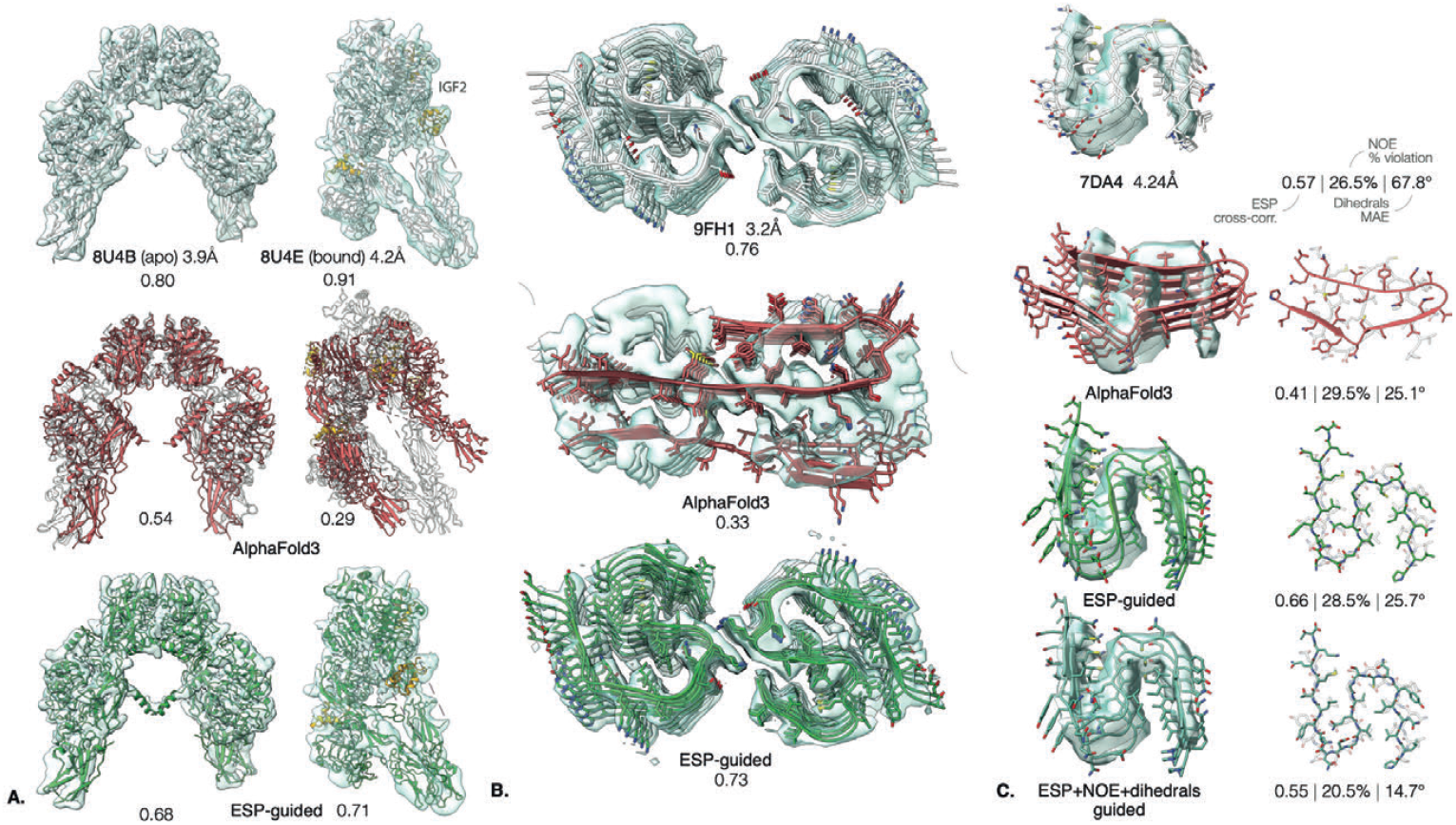
Multimeric structure reconstruction using cryoEM electrostatic potential (ESP) map guidance. (A) Reconstruction of insulin receptor IR-B in its symmetric apo state (left) and asymmetric conformation (right) with three bound IGF2 factors (yellow); and (B) reconstruction of an amyloid-beta fibril. AlphaFold3 (red) and ESP-guided (green) structures are shown alongside the deposited PDB models. Density cross-correlation values are indicated below each structure. (C) Reconstruction of a human RIPK3 amyloid fibril using unguided AlphaFold3 (red), ESP-guided AlphaFold3 (green) and ESP-guided AlphaFold3 with the addition of NMR chemical shift-inferred backbone dihedral angles ϕ, ψ (cyan). The addition of the dihedrals further improves agreement with both experimental observations. Cross-correlations, NOE constraint violations and dihedral errors are reported below each structure.

Fig. 6B shows the reconstruction of an amyloid-beta fibril (PDB 9FH1, EMDB 50436) formed by a nearly planar homodimer that further oligomerizes in the perpendicular direction forming a very long fiber structure. AlphaFold3 completely mispredicts the dimerization interface and the structure of the dimeric unit. ESP guidance produces accurate agreement with the density.

### Guiding AlphaFold3 jointly with cryoEM and NMR data

In many cases, structural biology approaches comprise data from several techniques. Our approach can integrate all the above sources of data, and more. We exemplify such a case here for the combination of cryoEM and NMR data, applied to amyloid fibrils. Fig. 6C shows the reconstruction of a RIPK3 human amyloid fibril (PDB 7DA4, EMDB 30622) for which solid-state NMR chemical shift-derived dihedral angles are also available (PDB 7DAC, BMRB 36392). In this case, AlphaFold3 predicts the global structure of the multimer relatively well within the resolution of the ESP map, and ESP-only guidance improves the density fit quality while producing a relatively poor fit of the backbone dihedrals and NOE constraints. Guiding jointly by ESP, NOE and dihedral angles significantly improves the local structure accuracy at a reasonable tradeoff of density fit (see SI Table 5 for quantitative analysis).

## Discussion

This work reframes AlphaFold3 as a powerful sequence-conditioned prior that can be steered by experimental evidence to yield small, testable conformational ensembles rather than a single consensus model. Casting structure determination as posterior inference makes it straightforward to compose likelihoods from heterogeneous modalities—inter-atomic distances, dihedral angles order parameters from NMR, electron density from X-ray crystallography, and electrostatic-potential maps from cryo-EM—and to direct sampling toward structures that better explain the data. In practice, guidance is injected as gradients during AF3’s reverse-diffusion trajectory and followed by relaxation and sparsifying ensemble selection, producing a reduced set of conformers that is both chemically plausible and data-faithful. The method’s ability to sample from the joint distribution of all atoms unlocks a richer picture of the protein’s conformational landscape that is partially lost in the traditional PDB representation of its structure as a mixture of Gaussian marginals of individual atoms (for X-ray data) or ensembles of conformers from NMR, which, as we show here, do not reflect dynamic amplitudes faithfully.

Currently, the field lacks human-intelligible visualization and machine-interpretable representations of the joint probability distributions that govern proteins and complexes. An intriguing question is whether Nature re-uses a restricted vocabulary of local distributional motifs—analogous to secondary structure—but defined over distributions of backbone/side-chain states, and whether there exist non-local distributional motifs of correlated motion and allostery, analogous to structural domains. The degree to which such motifs are imprinted in evolutionary signals (e.g., MSA features) will determine how far inductive models can progress from predicting static structure to predicting relative populations of states. The comparison to existing inductive ensemble generators such as BioEmu and AlphaFlow suggests that in the absence of experimental evidence, these models largely fail to produce ensembles in agreement with the experiment.

Methodologically, three features are central in this study. First, ensembles are jointly guided during diffusion, which preserves diversity where the data admit multiple explanations while avoiding trivial replication of a single conformation; subsequent relaxation ensures chemical plausibility; and a matching-pursuit–like pruning step yields a compact set that still explains the observations. Second, the framework naturally supports “local-to-global” strategies—targeting problematic segments (e.g., loops) and then iterating to the whole chain—mirroring how crystallographers and NMR spectroscopists often work. Third, the computational footprint is small (minutes per target in typical cases, which is likely to be further optimizable), making the method practical as an interactive assistant within existing pipelines rather than a replacement for refinement or simulation. For the same reason, the tool enables large-scale re-analysis of legacy datasets (“PDB redo”) across single modalities and their combinations.

Several caveats warrant emphasis. The reported ensembles are explanatory with respect to the provided data and the AF3 prior; they are not calibrated thermodynamic populations. Without additional constraints, mixture weights should not be over-interpreted as equilibrium populations. Future work could link pruning weights to maximum-entropy or evidence-based population estimates, or incorporate sparse thermodynamic measurements (e.g., ΔG) to at least partially calibrate populations. Guidance strength and gradient handling are important guardrails to avert models being pulled into noise, especially in low-resolution EM or poorly phased crystallographic maps. While in practice ensemble pruning and cross-validation techniques mitigate overfitting, a more principled statistically robust procedure is needed to disentangle true thermodynamic entropy from epistemic uncertainty. Finally, the present implementation focuses primarily on proteins; treatment of ligands, metals, PTMs, structural waters, and complex symmetries remains limited when the only evidence is a density-like signal. A more faithful modeling of ligands will also allow disentangling between native thermodynamic and compositional conformational heterogeneity.

These limitations notwithstanding, the posterior-sampling view opens practical routes to improve mainstream structural workflows. In crystallography, experiment-guided AlphaFold3 can propose altlocs and occupancies during model building, repair loops with weak electron density, and accelerate convergence by offering a small set of candidate conformers that already satisfy the map. In NMR, it enables rapid, ensemble-aware interpretation of NOESY spectra and relaxation data without days of MD, and can suggest mutations or ligands that stabilize minor predicted states for validation. In cryo-EM, it provides a principled way to inject atomic detail into flexible regions and to couple map-based guidance with dihedral or secondary-structure information where local resolution is limited. We anticipate that experiment-guided generative models will make ensemble-centric modeling routine in crystallography, NMR, and cryo-EM, with near-term impact on ligand discovery, variant interpretation, and the design of experiments that stabilize or reveal otherwise cryptic states.

## Data and Code Availability

All structures and metrics reported in this paper are available at https://doi.org/10.7910/DVN/PLYUHN

The code is available at https://doi.org/10.5281/zenodo.17307005

## Acknowledgments

AM acknowledges the financial support of the Helmsley Fellowships Program for Sustainability and Health. AMB and PS are supported by the ISTA IPC grant *Generative Protein NMR*. SV was supported in part by funding from the Eric and Wendy Schmidt Center at the Broad Institute of MIT and Harvard.

## Supplementary Information

## 1 Supplementary Figures and Tables

**SI Figure 1:**
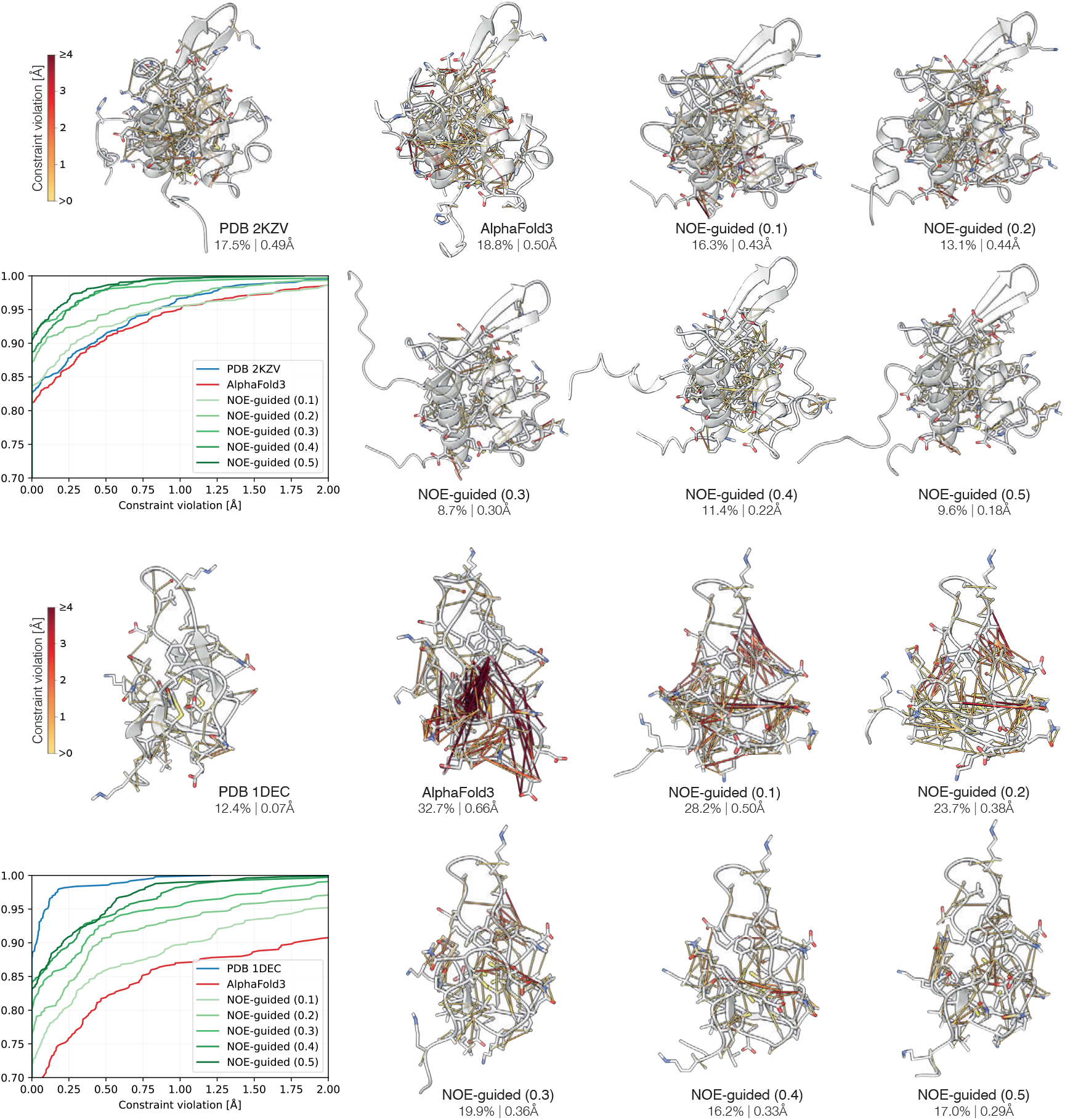
Guidance of `2KZV` (top) and `1DEC` (bottom) structure by NOE pairwise distance bounds with varying data term strength. Visualized are the pairwise distance constraint violations for the PDB ensemble, unguided AlphaFold3, and NOE-guided predictions with varying data term strength indicated in parenthesis. The fraction of the violated constraints and their median violation are reported below each structure. A single best-fitting structure from the ensemble is shown for clarity. Plots show cumulative distribution of constraint violations.

**SI Figure 2:**
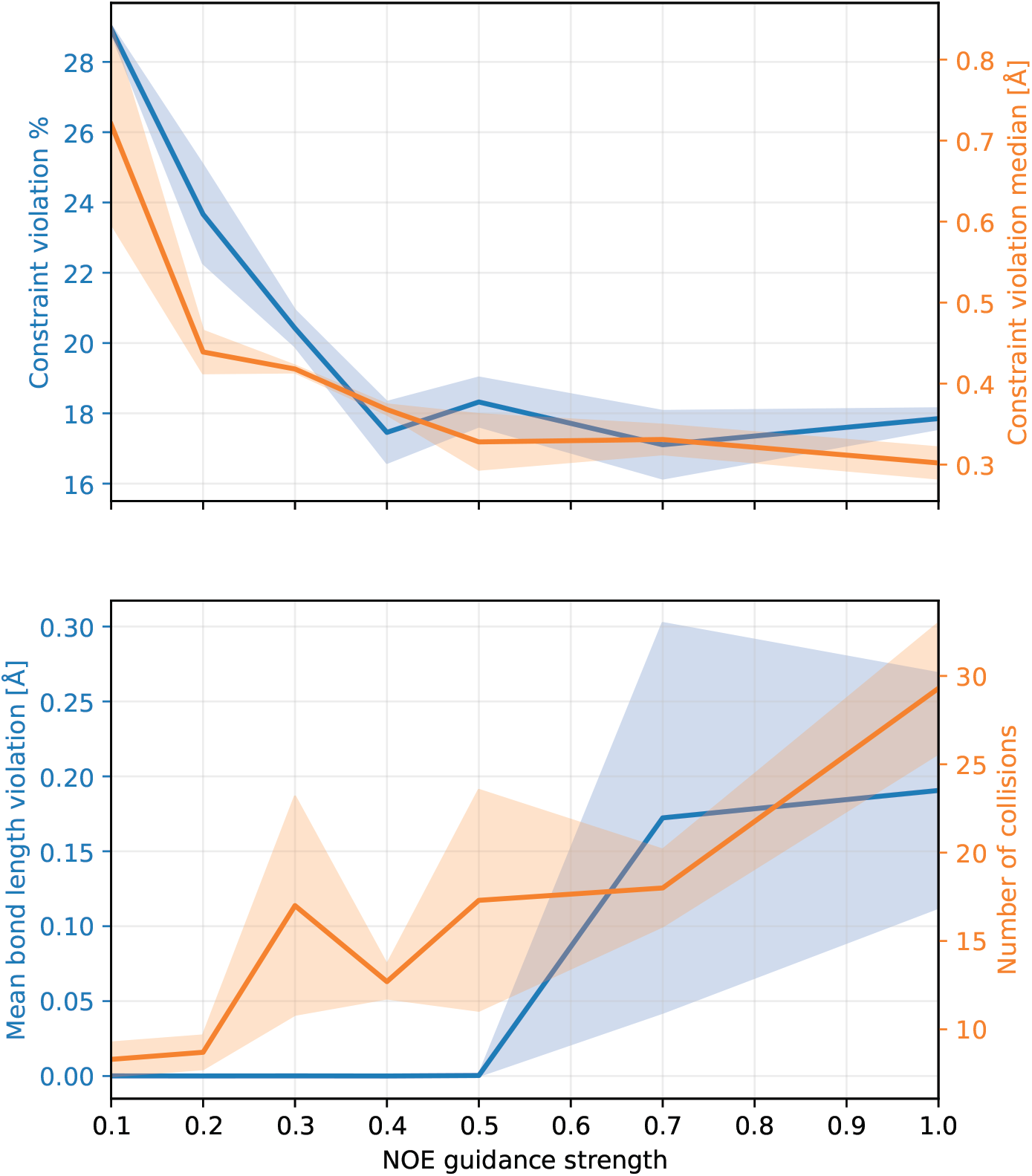
The tradeoff between agreement with experimental data and the feasibility of the generated structures demonstrated on NOE guidance of `1DEC`. Plotted are the distance constraint violations (percentage and median), bond length deviation from the nominal values, and number of atom clashes. Solid lines indicate averages on 3 runs; areas depict standard deviations. In our experiments, we set the guidance strength to 0.4 unless indicated otherwise.

**SI Figure 3:**
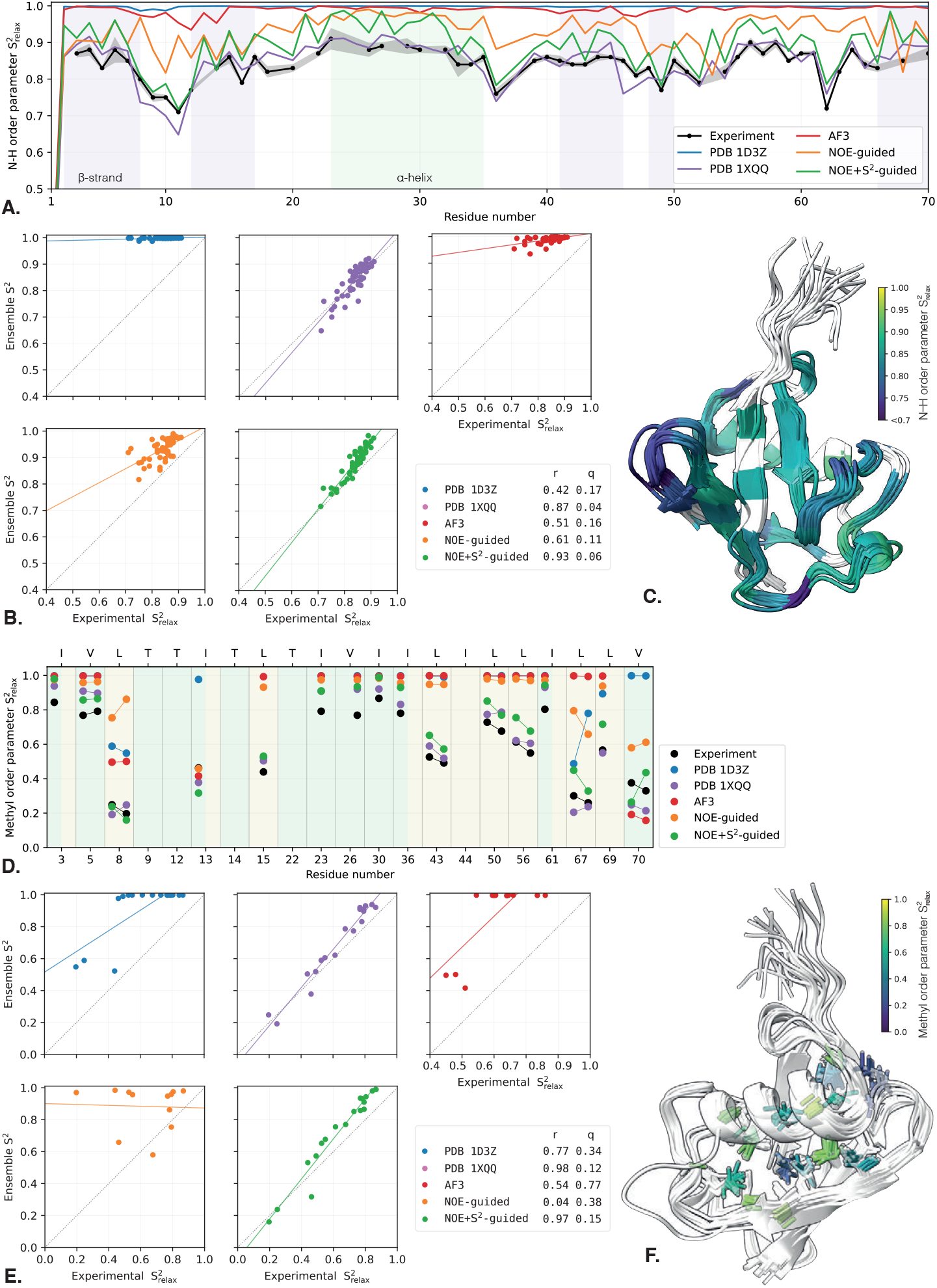
Guidance of ubiquitin structure with the combination of NOE pairwise distance bounds with amide (top) and methyl (bottom) order parameters obtained from relaxation data (ps-ns motion). (A) Experimentally measured amide order parameter *S*^2^ from [22] (black) vs. order parameter calculated from the ensembles of different methods. (B) The same data visualized as a scatter plot with correlation (*r*, higher is better) and normalized fitting error (*q*, lower is better) factors. (C) Visualization of the disorder parameter on the NOE+*S*^2^-guided ensemble. Lower values mean bigger motion. (D) Experimentally measured methyl order parameter *S*^2^ from [21] (black) vs. order parameter calculated from the ensembles of different methods. C_*γ*_ and C_*δ*_ groups are indicated in green and yellow, respectively. For instances with two data points, these refer to the two methyl groups of Val (*γ*_1_, *γ*_2_), Leu (*δ*_1_, *δ*_2_) and Ile (*γ*_2_, *δ*_1_). (E) Same data visualized as scatter plots. (F) Visualization of the disorder parameter on the NOE+*S*^2^-guided ensemble with relevant bonds colored by their *S*^2^ values.

**SI Figure 4:**
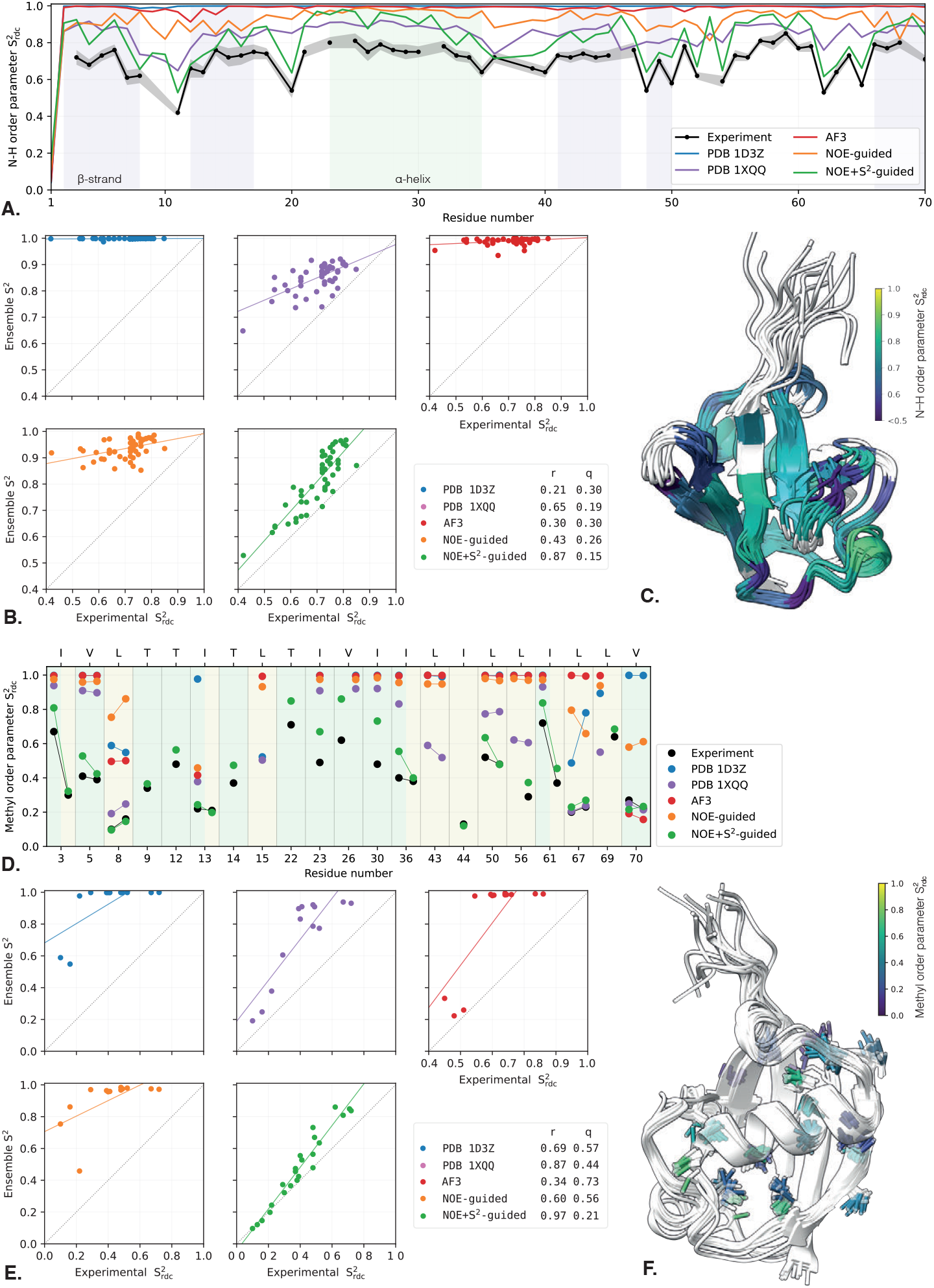
Guidance of ubiquitin structure with the combination of NOE pairwise distance bounds with amide (top) and methyl (bottom) order parameters obtained from residual dipolar coupling data (*µ*s-ms motion). (A) Experimentally measured amide order parameter *S*^2^ from [31] (black) vs. order parameter calculated from the ensembles of different methods. (B) The same data visualized as a scatter plot with correlation (*r*, higher is better) and normalized fitting error (*q*, lower is better) factors. (C) Visualization of the disorder parameter on the NOE+*S*^2^-guided ensemble. Lower values mean bigger motion. (D) Experimentally measured methyl order parameter *S*^2^ from [9] (black) vs. order parameter calculated from the ensembles of different methods. C_*γ*_ and C_*δ*_ groups are indicated in green and yellow, respectively. For instances with two data points, these refer to the two methyl groups of Val (*γ*_1_, *γ*_2_), Leu (*δ*_1_, *δ*_2_) and Ile (*γ*_2_, *δ*_1_). (E) Same data visualized as scatter plots. (F) Visualization of the disorder parameter on the NOE+*S*^2^-guided ensemble with relevant bonds colored by their *S*^2^ values.

**SI Figure 5:**
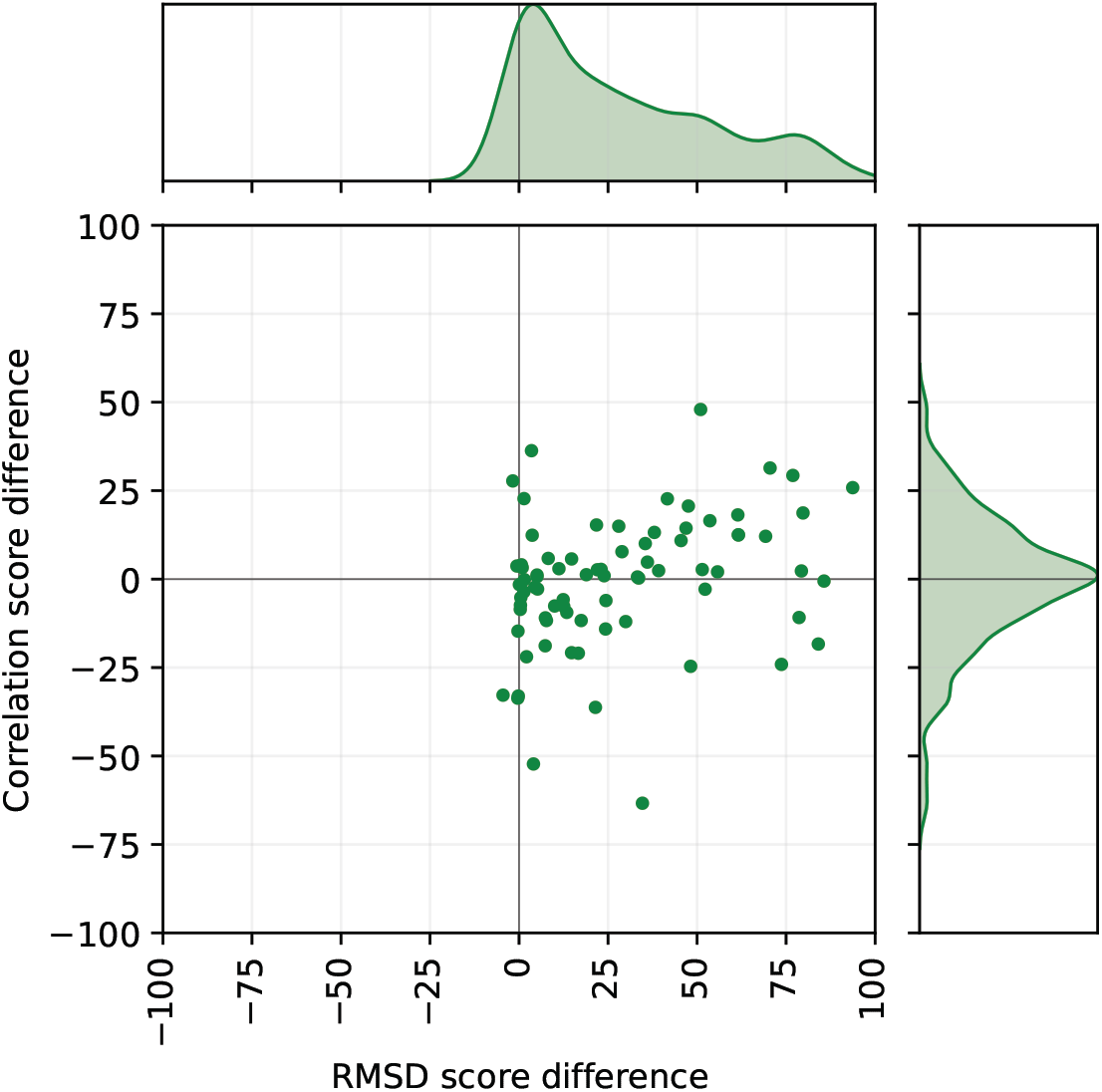
ANSURR [11] measures how well a structure’s residue-by-residue rigidity profile agrees with that inferred from NMR chemical shifts: the correlation score captures similarity of the pattern (where flexible/rigid regions occur), while the RMSD score captures agreement in the magnitude of rigidity/flexibility across residues. Scores are normalized from 0 to 100 with higher numbers indicating better agreement. Depicted is the paired difference in the two ANSURR scores between the NOE-guided ensembles and the corresponding PDB structures. Each point corresponds to one of the evaluated 77 structures for which the median shift was above 80%. Marginal plots show kernel density estimates of each score individually. Our guidance method consistently improves the RMSD score while maintaining comparable correlation scores.

**SI Figure 6:**
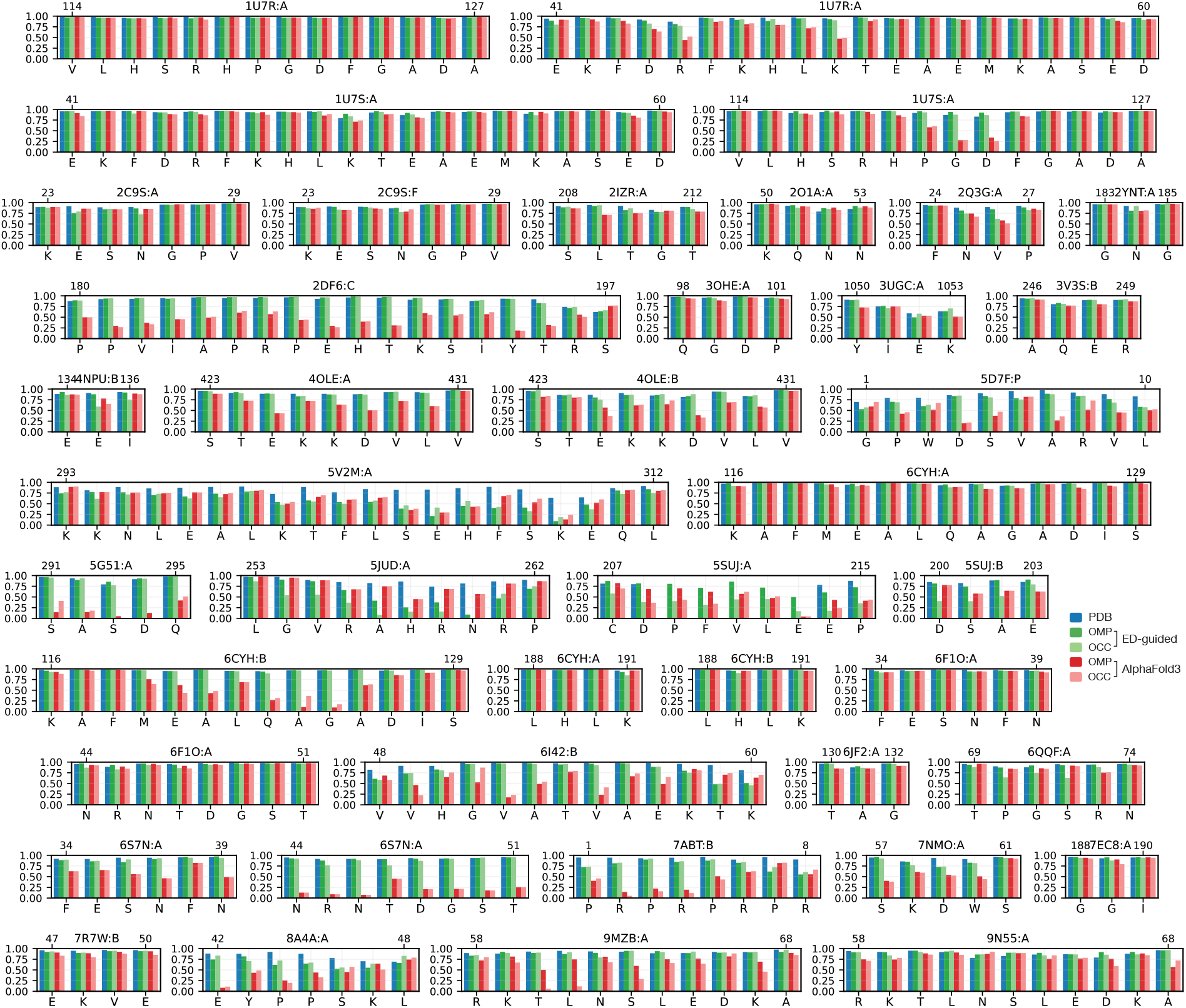
A summary of per-residue real-space correlation coefficients (RSCC) for the evaluated X-ray crystallographic structures reported in Table 2. Color-coded bars represent the RSCC of the deposited PDB structure (blue), electron density-guided AlphaFold3 with OMP (dark green) and occupancy optimization (light green) ensemble pruning, and unguided AlphaFold3 with OMP (dark red) and occupancy optimization (light red) ensemble pruning.

**SI Figure 7:**
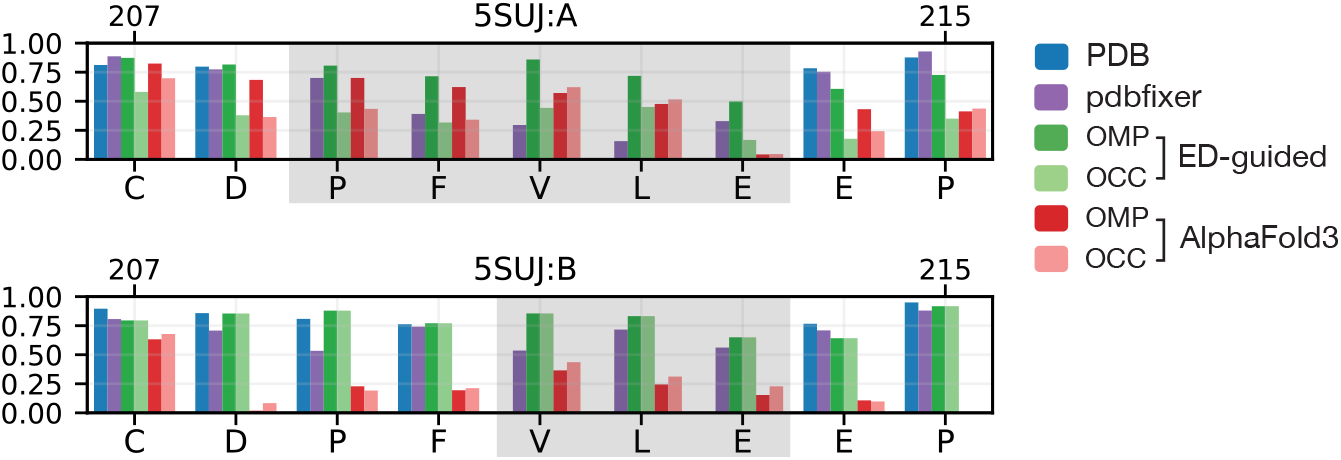
Per-residue real-space correlation coefficients (RSCC) for the `5SUJ` X-ray crystallographic structure with unmodelled density (highlighted in grey). Color-coded bars represent the RSCC of the deposited PDB structure (blue), completed structure with pdbfixer (purple), electron density-guided AlphaFold3 with OMP (dark green) and occupancy optimization (light green) ensemble pruning, and unguided AlphaFold3 with OMP (dark red) and occupancy optimization (light red) ensemble pruning.

**SI Figure 8:**
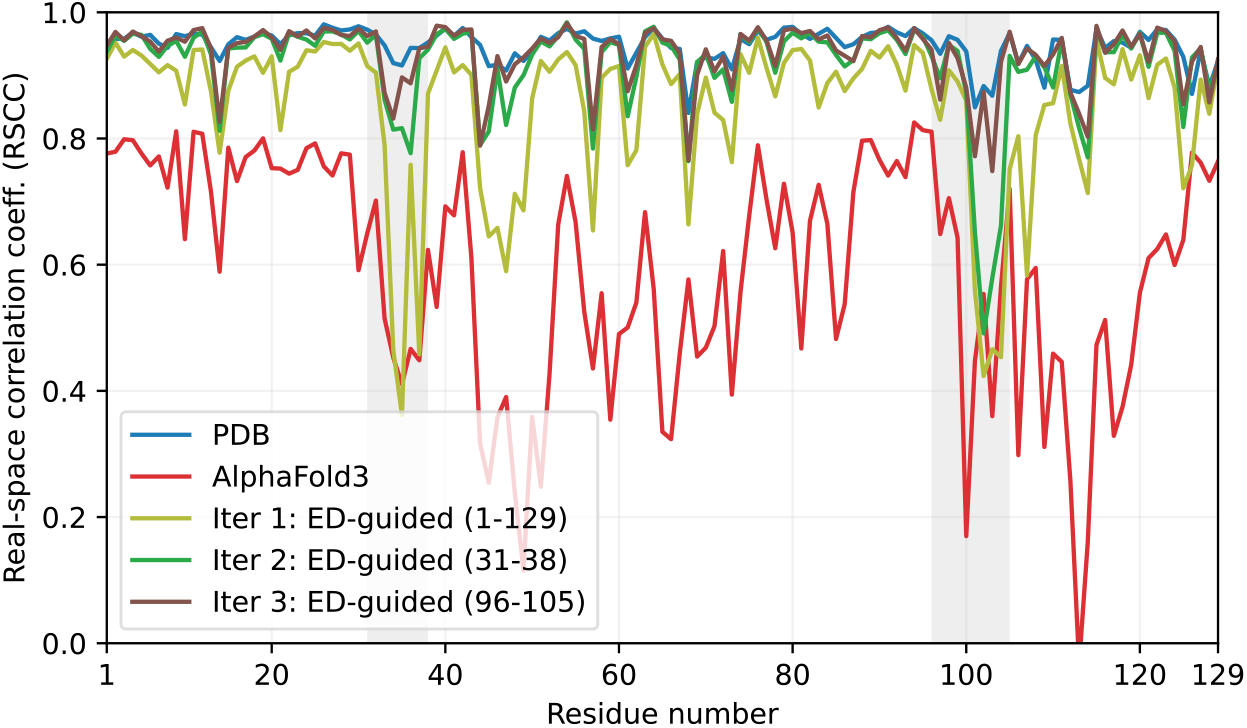
Per-residue real-space correlation coefficients (RSCC) for the `6S7N` X-ray crystallographic structure modeled with electron-density (ED) guided AlphaFold3 iterated with real-space refinement. Color-coded lines represent the RSCC of the deposited PDB structure (blue), unguided AlphaFold3 prediction (red), first iteration in which the entire structure was ED-guided (olive), followed by phenix real-space refinement and a second iteration of partial structure guidance for residues 1 − 38 (green), followed by phenix refinement and a third iteration of partial structure guidance for residues 96 − 105 (brown). Further iterations with partial guidance in worst-performing regions show further albeit diminishing improvement in RSCC.

**SI Figure 9:**
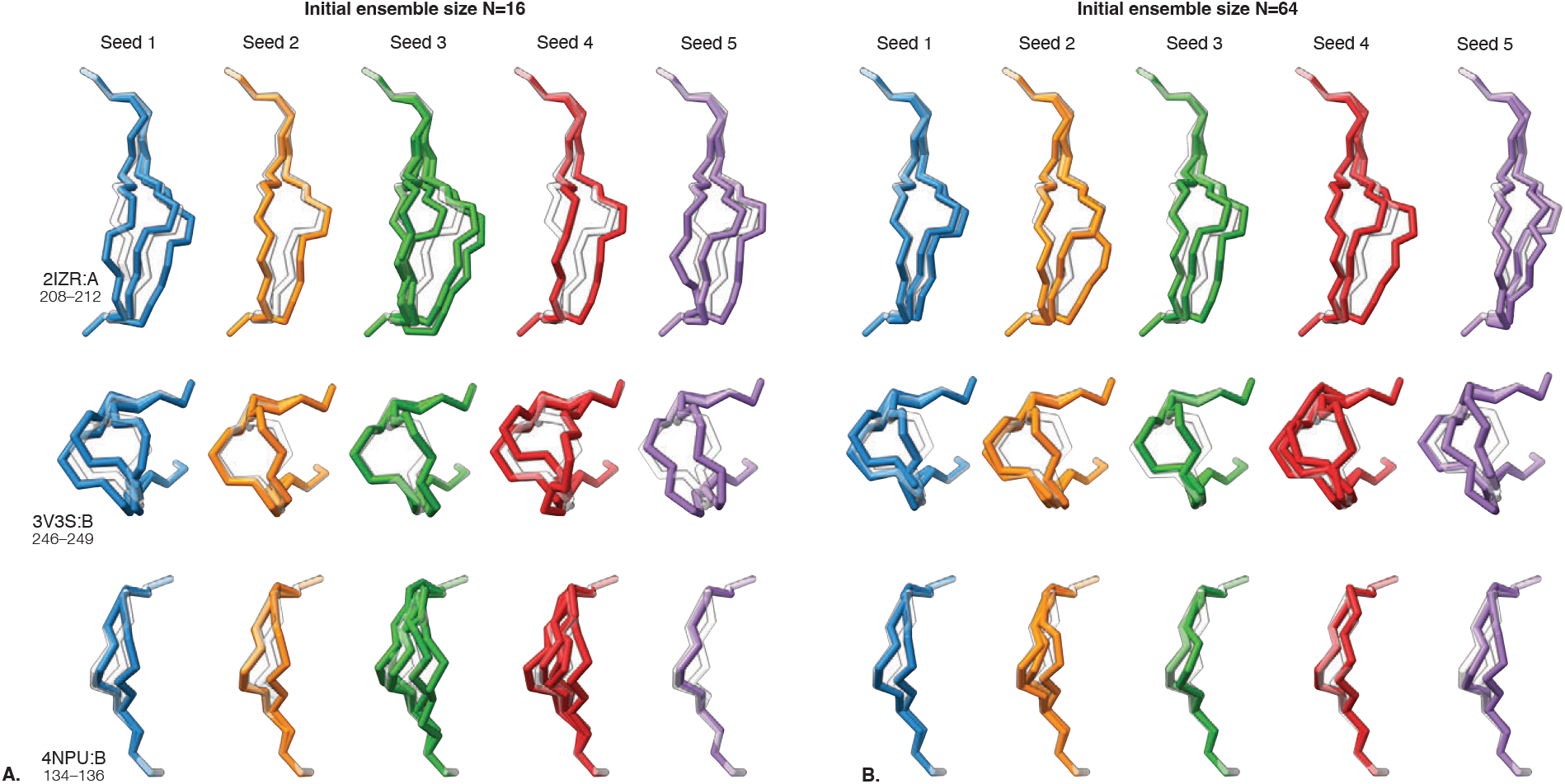
Repeatability of generated ensembles with five random seeds using initial ensemble sizes of 16 (A) and 64 (B) members. The generated backbones are overlaid on top of the corresponding PDB models depicted in white.

**SI Figure 10:**
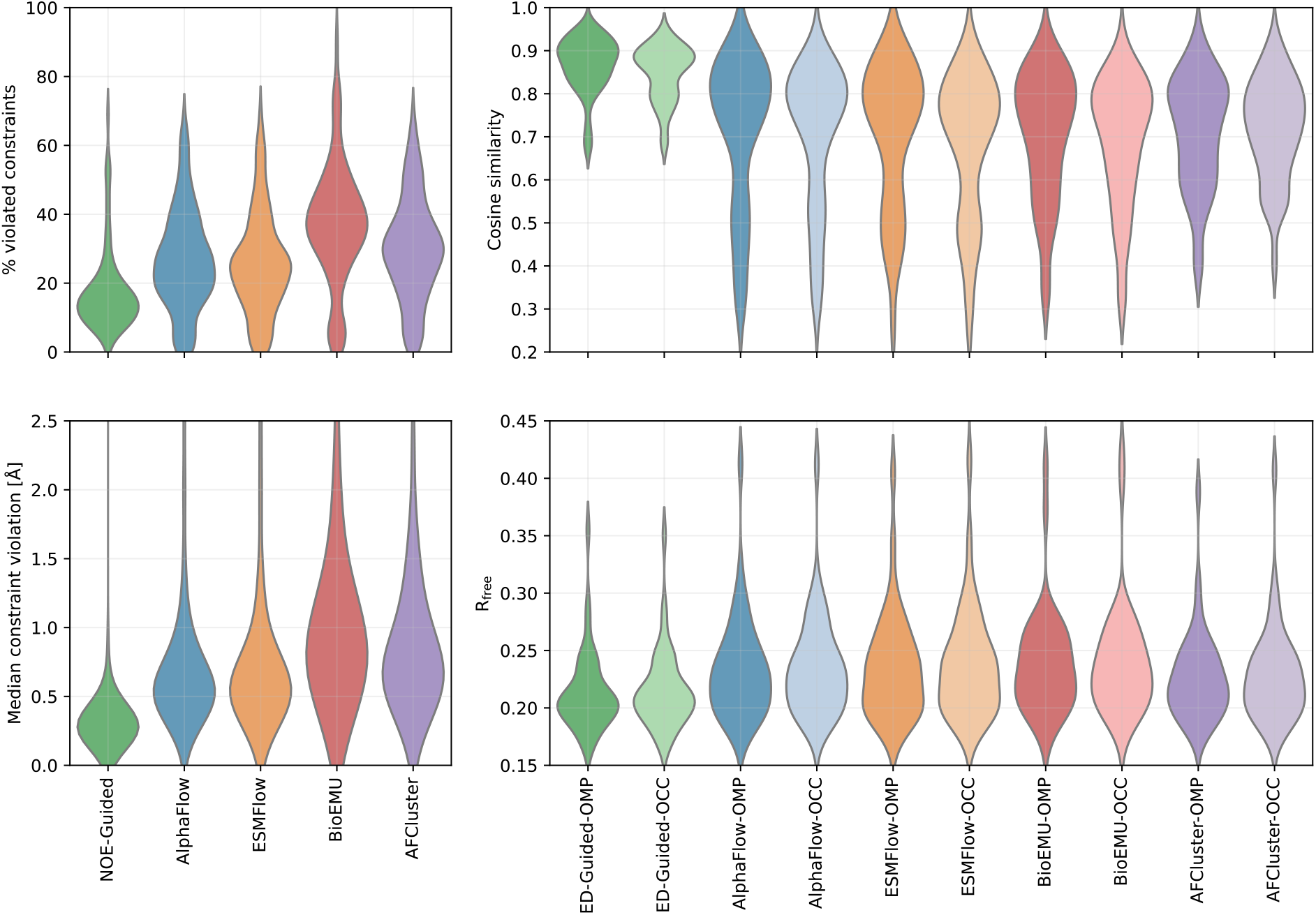
Left: Distribution of % of violated distance constraints (top) and median violation (bottom) for NOE-guided AlphaFold3 and various ensemble generation models. Right: distribution of F_o_ and F_c_ cosine similarity (top) and R_free_ (bottom) for ED-guided AlphaFold3 and the same ensemble generators. Shown are OMP and occupancy optimization (OCC) ensemble prunning for all models.

**SI Figure 11:**
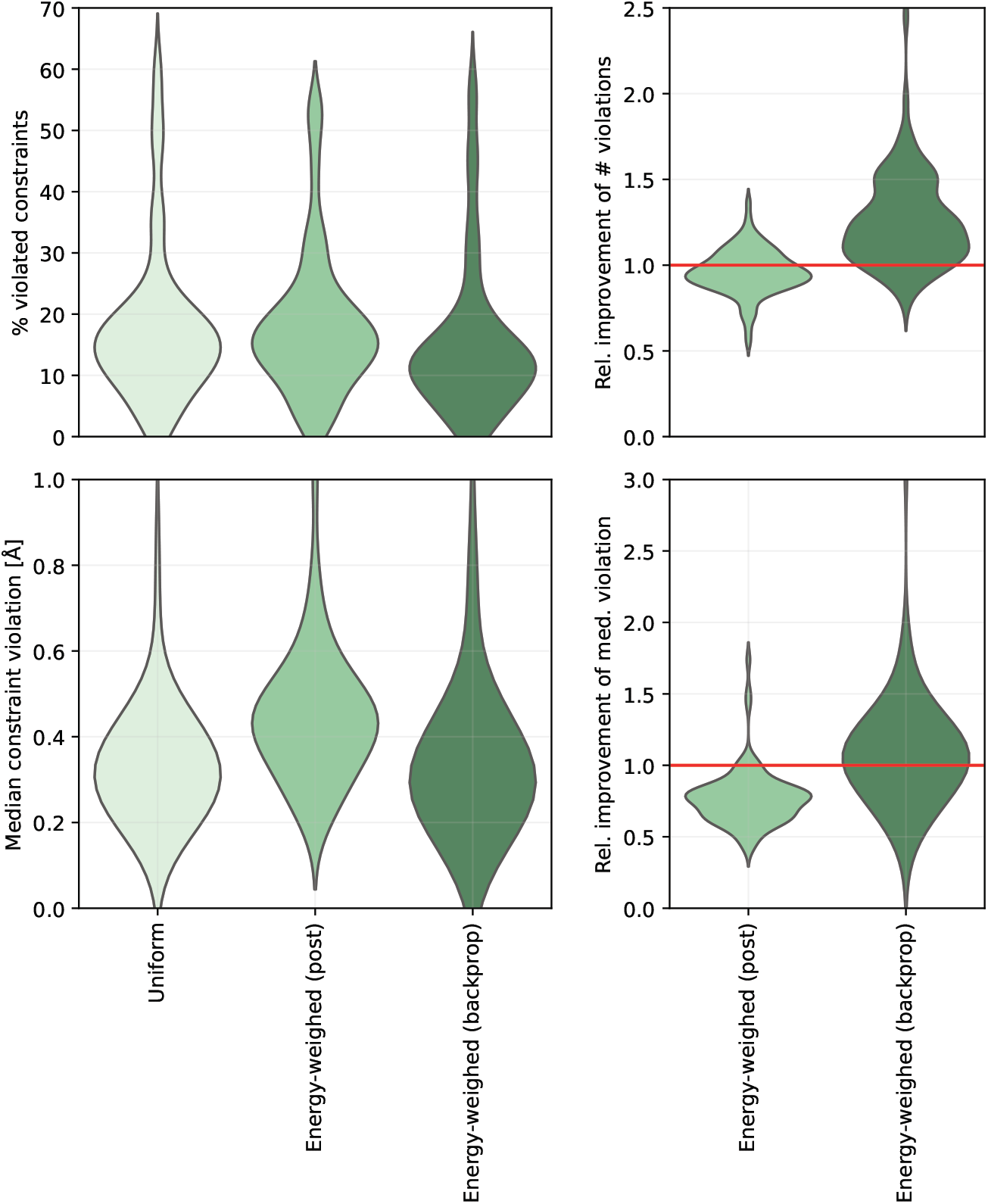
Left: Distribution of % of violated distance constraints (top) and median violation (bottom) for NOE-guided AlphaFold3 with various ensemble weighing techniques: uniform, energy-weighed at post-processing, and energy-weighed with full backpropagation through the force field during guidance. Right: the relative improvement of these metrics compared to the uniform weighing. This figure complements Figure 5A in the main text.

**SI Table 1:**
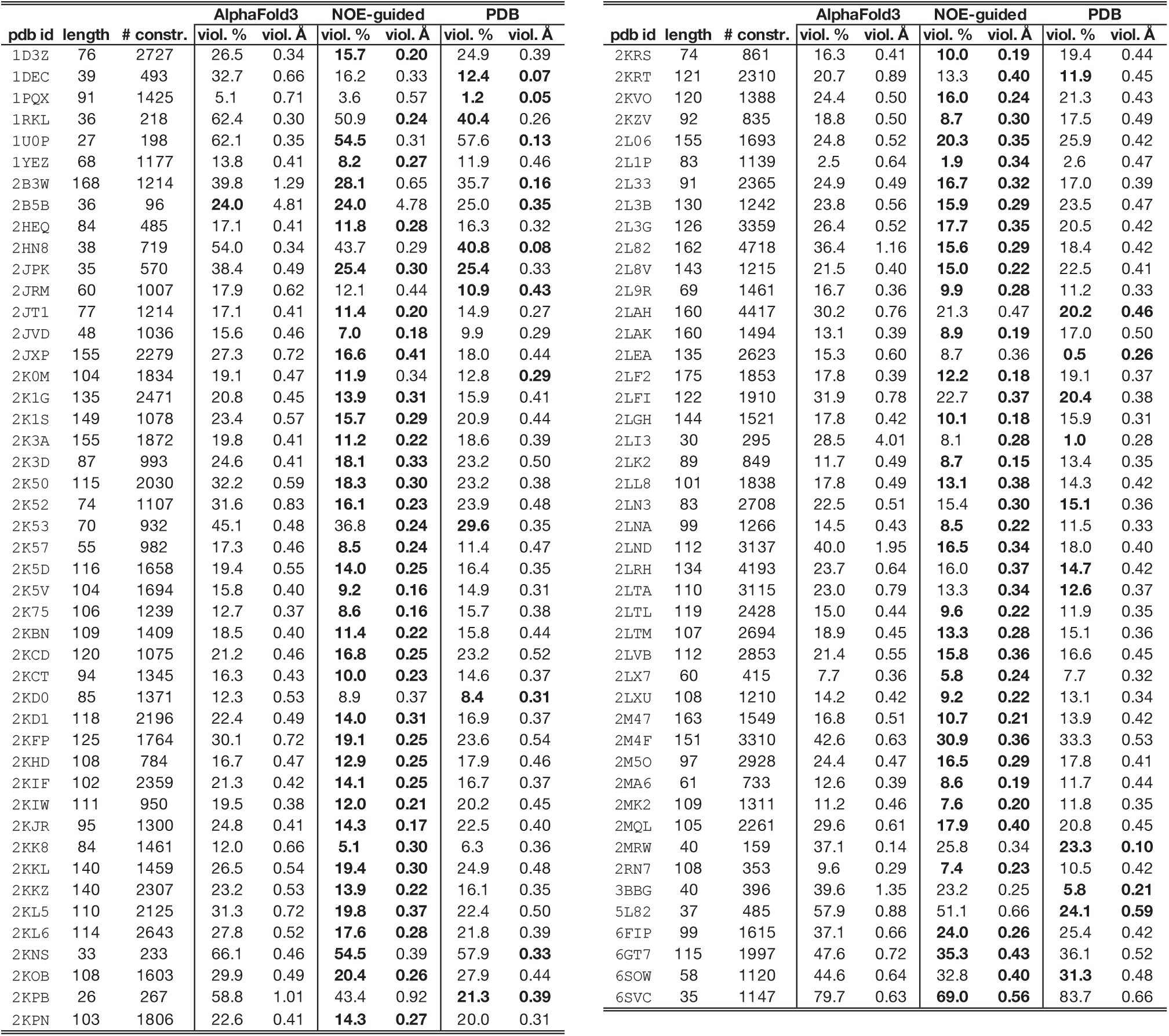
Summary of quantitative performance indicators of the evaluated NMR structures. Reported are the number of NOE constraints, percentage of constraint violation and median magnitude of the violated constraints for unguided and NOE-guided AlphaFold3 and the corresponding deposited PDB structures.

**SI Table 2:**
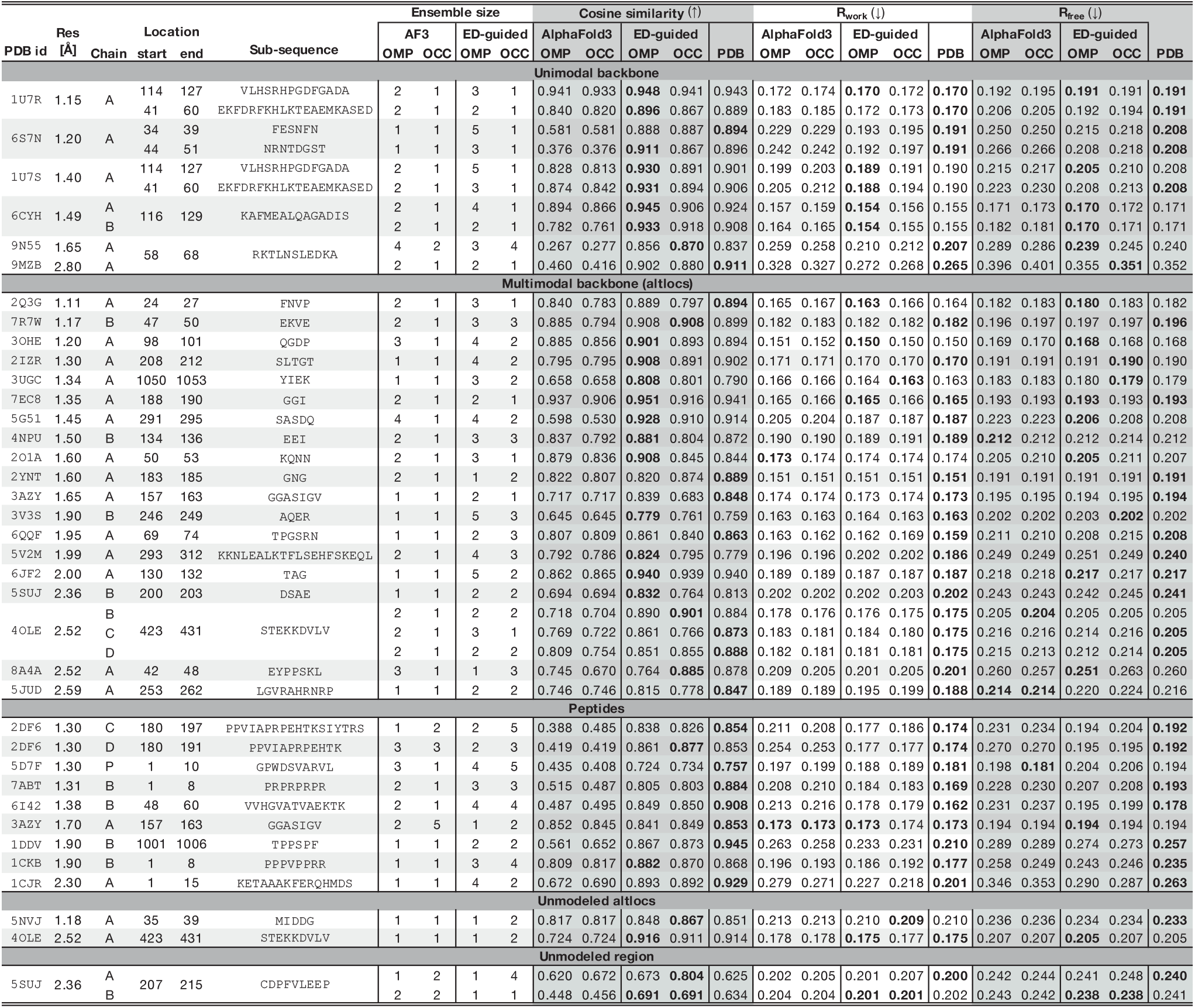
Summary of quantitative performance indicators of the evaluated X-ray crystallographic structures. Reported are the structure resolution, the location of the generated region, cosine similarity as the real-space metric and the *R*-factors (*R*_work_ and *R*_free_) as the Fourier-space metrics for unguided and electron density-guided AlphaFold3 and the corresponding deposited PDB structures. Ensemble sizes produced by OMP and occupancy optimization (OCC) pruning are reported in the latter cases.

**SI Table 3:**
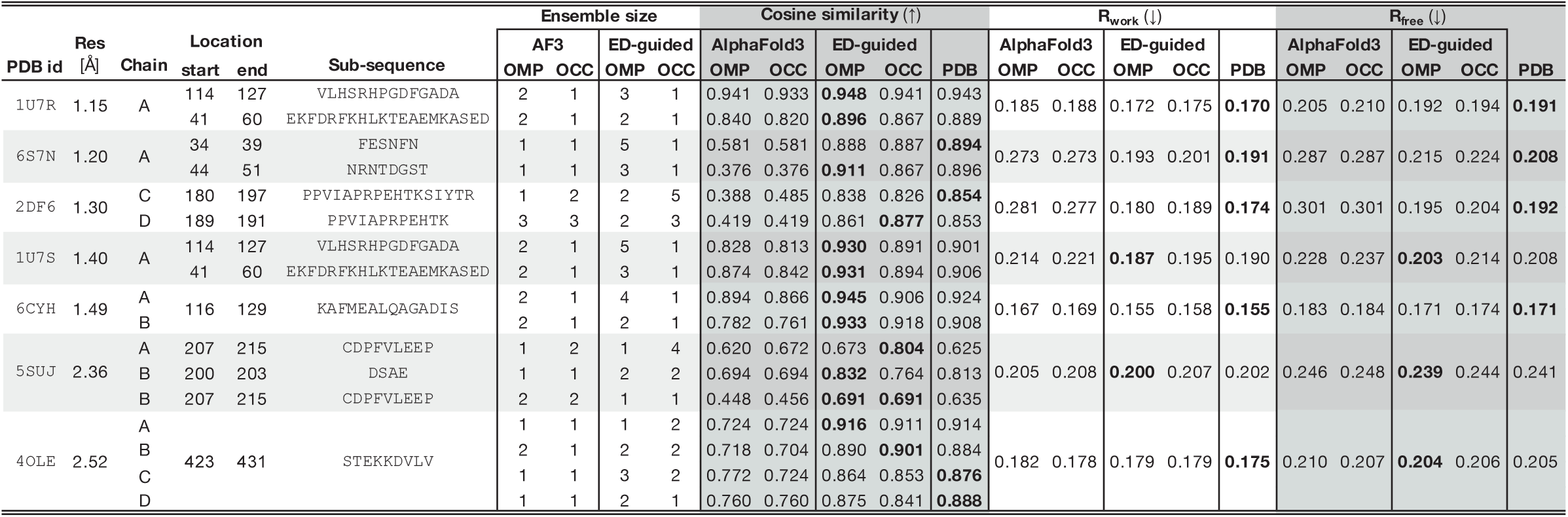
A subset of structures from Table 2 in which several locations in the same structure were merged and jointly refined.

**SI Table 4:**
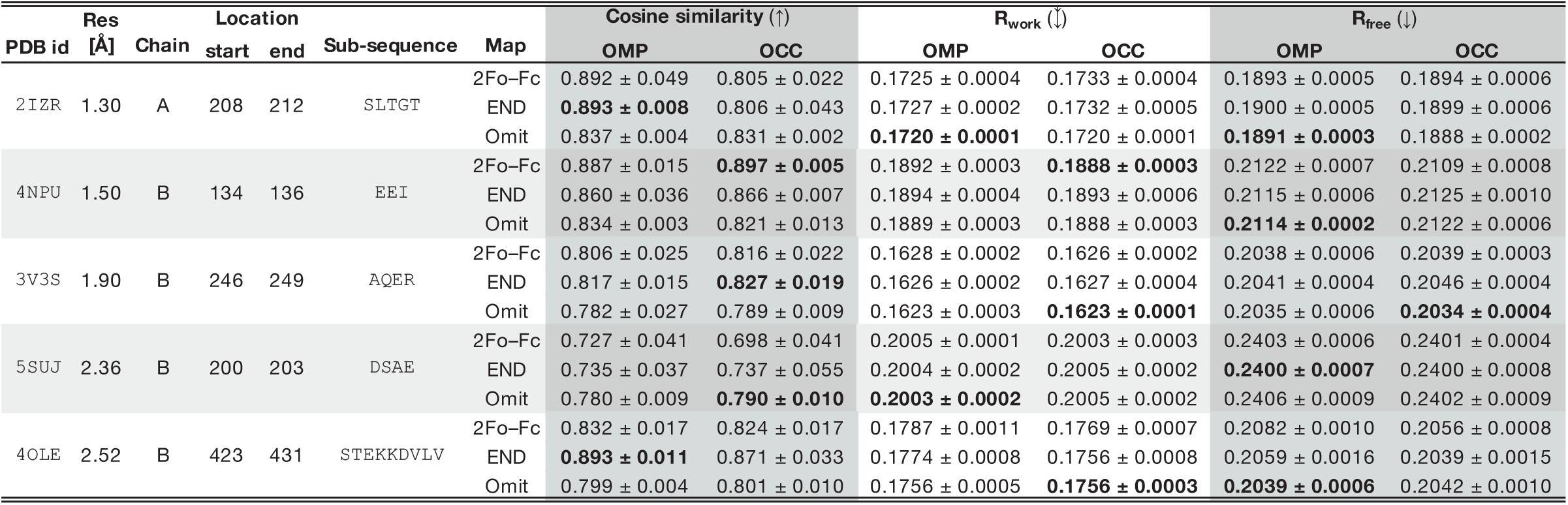
ED-guided AlphaFold3 reconstruction on a subset of structures from Table 2. Shown are average metrics ± standard deviation over 5 independent runs with different random seeds using three types of inputs: 2*F*_o_ − *F*_c_, END, and omit maps. Note that cosine similarities, being real-space metrics, are not directly comparable across different map types.

| PDB id | Res<br>[Å] | Chain | Location |  | Sub-sequence | Map | Cosine similarity (†) |  |  | R<sub>work</sub> (‡) |  |  | R<sub>free</sub> (‡) |  |
| --- | --- | --- | --- | --- | --- | --- | --- | --- | --- | --- | --- | --- | --- | --- |
|  |  |  | start | end |  |  | OMP | OCC |  | OMP | OCC |  | OMP | OCC |
| 2IZR | 1.30 | A | 208 | 212 | SLTGT | 2Fo-Fc | 0.892 ± 0.049 | 0.805 ± 0.022 |  | 0.1725 ± 0.0004 | 0.1733 ± 0.0004 |  | 0.1893 ± 0.0005 | 0.1894 ± 0.0006 |
|  |  |  |  |  |  | END | **0.893 ± 0.008** | 0.806 ± 0.043 |  | 0.1727 ± 0.0002 | 0.1732 ± 0.0005 |  | 0.1900 ± 0.0005 | 0.1899 ± 0.0006 |
|  |  |  |  |  |  | Omit | 0.837 ± 0.004 | 0.831 ± 0.002 |  | **0.1720 ± 0.0001** | 0.1720 ± 0.0001 |  | **0.1891 ± 0.0003** | 0.1888 ± 0.0002 |
| 4NPU | 1.50 | B | 134 | 136 | EEI | 2Fo-Fc | 0.887 ± 0.015 | **0.897 ± 0.005** |  | 0.1892 ± 0.0003 | **0.1888 ± 0.0003** |  | 0.2122 ± 0.0007 | 0.2109 ± 0.0008 |
|  |  |  |  |  |  | END | 0.860 ± 0.036 | 0.866 ± 0.007 |  | 0.1894 ± 0.0004 | 0.1893 ± 0.0006 |  | 0.2115 ± 0.0006 | 0.2125 ± 0.0010 |
|  |  |  |  |  |  | Omit | 0.834 ± 0.003 | 0.821 ± 0.013 |  | 0.1889 ± 0.0003 | 0.1888 ± 0.0003 |  | **0.2114 ± 0.0002** | 0.2122 ± 0.0006 |
| 3V3S | 1.90 | B | 246 | 249 | AQER | 2Fo-Fc | 0.806 ± 0.025 | 0.816 ± 0.022 |  | 0.1628 ± 0.0002 | 0.1626 ± 0.0002 |  | 0.2038 ± 0.0006 | 0.2039 ± 0.0003 |
|  |  |  |  |  |  | END | 0.817 ± 0.015 | **0.827 ± 0.019** |  | 0.1626 ± 0.0002 | 0.1627 ± 0.0004 |  | 0.2041 ± 0.0004 | 0.2046 ± 0.0004 |
|  |  |  |  |  |  | Omit | 0.782 ± 0.027 | 0.789 ± 0.009 |  | 0.1623 ± 0.0003 | **0.1623 ± 0.0001** |  | 0.2035 ± 0.0006 | **0.2034 ± 0.0004** |
| 5SUJ | 2.36 | B | 200 | 203 | DSAE | 2Fo-Fc | 0.727 ± 0.041 | 0.698 ± 0.041 |  | 0.2005 ± 0.0001 | 0.2003 ± 0.0003 |  | 0.2403 ± 0.0006 | 0.2401 ± 0.0004 |
|  |  |  |  |  |  | END | 0.735 ± 0.037 | 0.737 ± 0.055 |  | 0.2004 ± 0.0002 | 0.2005 ± 0.0002 |  | **0.2400 ± 0.0007** | 0.2400 ± 0.0008 |
|  |  |  |  |  |  | Omit | 0.780 ± 0.009 | **0.790 ± 0.010** |  | **0.2003 ± 0.0002** | 0.2005 ± 0.0002 |  | 0.2406 ± 0.0009 | 0.2402 ± 0.0009 |
| 4OLE | 2.52 | B | 423 | 431 | STEKKDVLV | 2Fo-Fc | 0.832 ± 0.017 | 0.824 ± 0.017 |  | 0.1787 ± 0.0011 | 0.1769 ± 0.0007 |  | 0.2082 ± 0.0010 | 0.2056 ± 0.0008 |
|  |  |  |  |  |  | END | **0.893 ± 0.011** | 0.871 ± 0.033 |  | 0.1774 ± 0.0008 | 0.1756 ± 0.0008 |  | 0.2059 ± 0.0016 | 0.2039 ± 0.0015 |
|  |  |  |  |  |  | Omit | 0.799 ± 0.004 | 0.801 ± 0.010 |  | 0.1756 ± 0.0005 | **0.1756 ± 0.0003** |  | **0.2039 ± 0.0006** | 0.2042 ± 0.0010 |

**SI Table 5:**
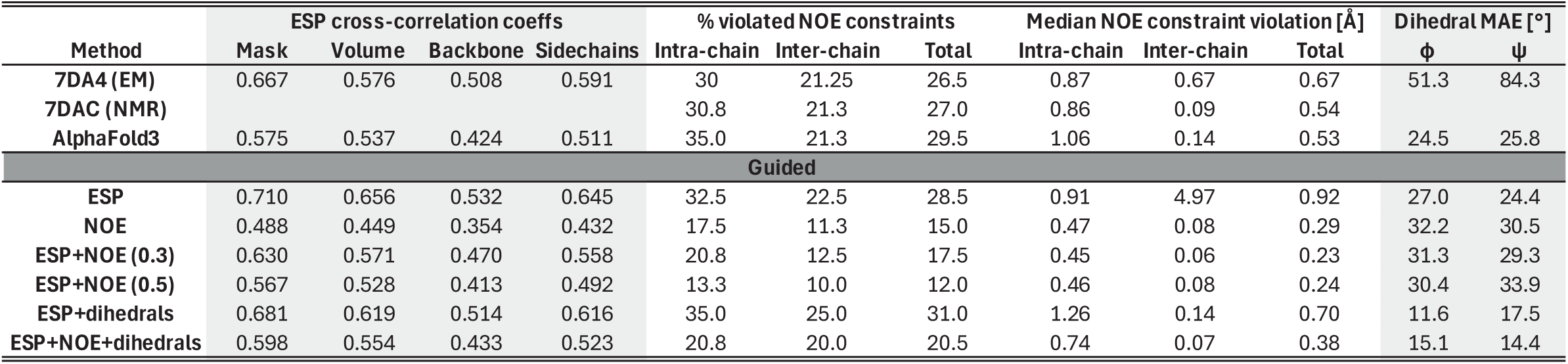
Guidance of `7DA4` with combinations of ESP, NOE and dihedral angles. Reported are cross-correlations with the ESP image, NOE constraint violations (within each chain and between the chains), and mean absolute disagreement with the chemical shift-derived backbone dihedral angles.

**SI Table 6:**
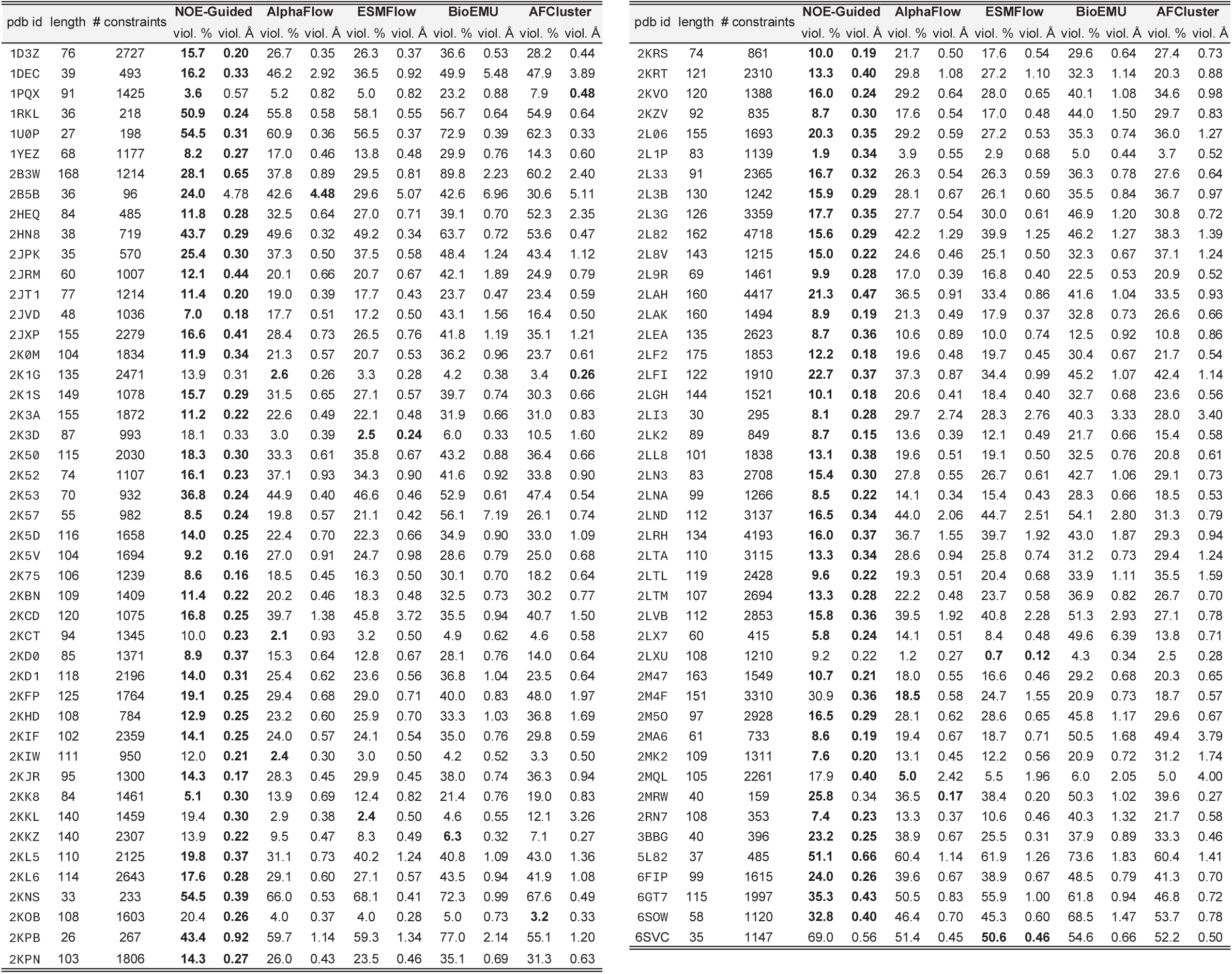
Evaluation of different ensemble generation models on NMR structures from Table 1.

**SI Table 7:**
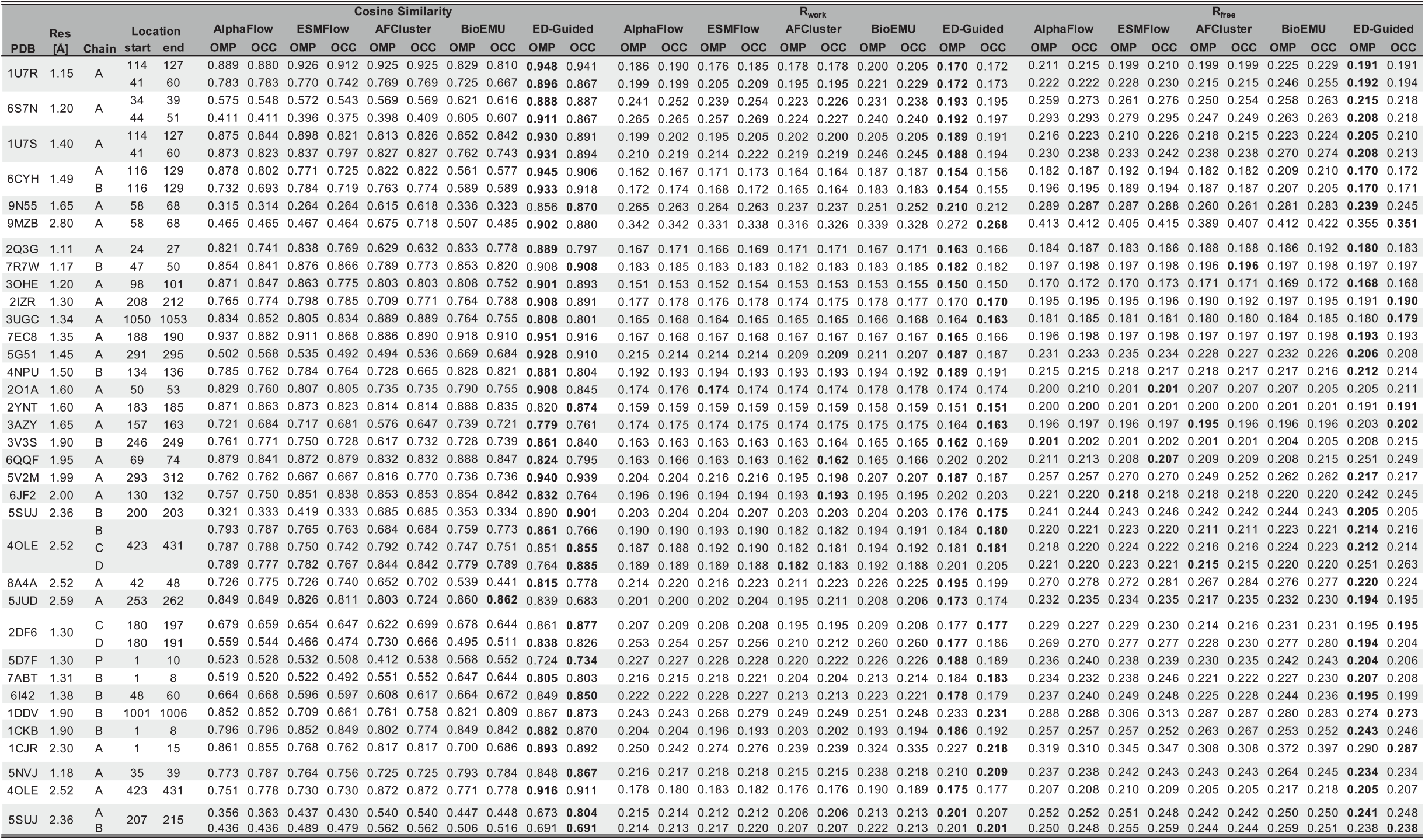
Evaluation of different ensemble generation models on X-ray structures from Table 2.

**SI Table 8:**
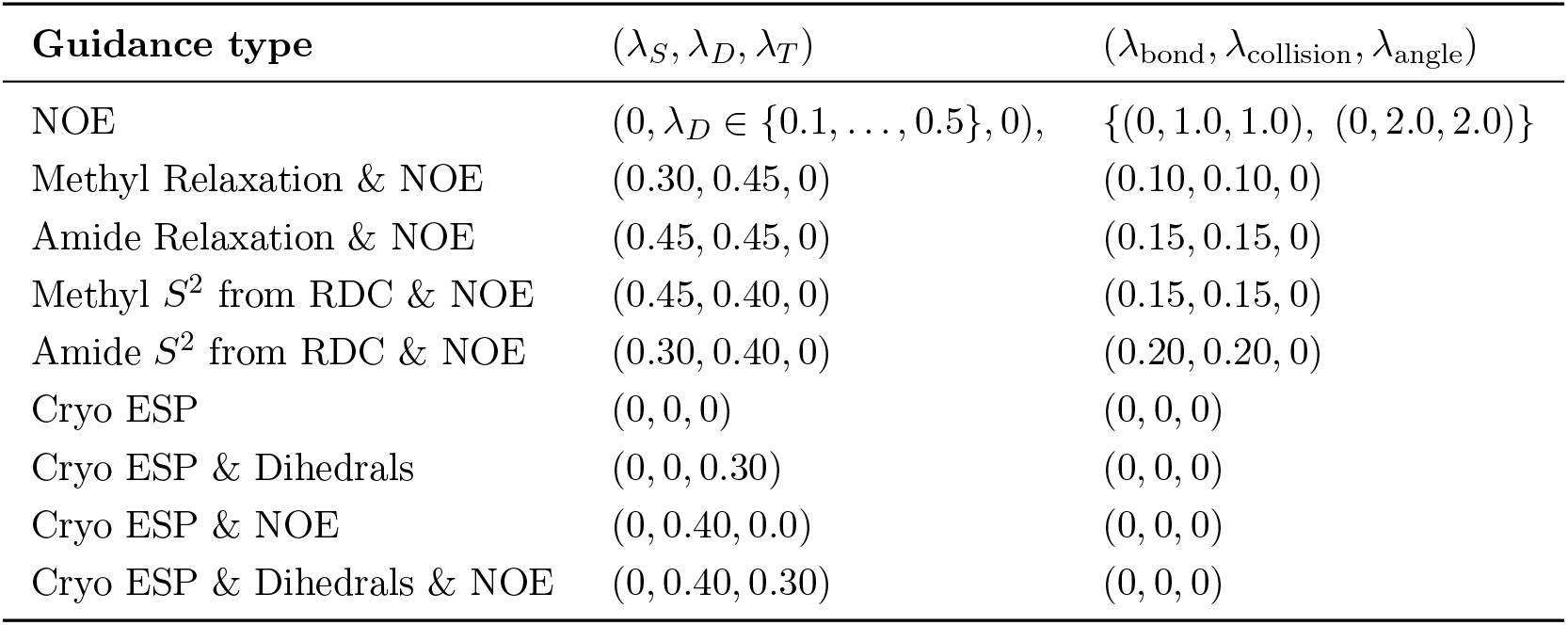
Scaling factors for NMR-guided ensembles.

## 2 Supplementary Methods

### 2.1 Models, resources, and packages

For all experiments, we used Protenix [34] – an open-sourced PyTorch re-implementation of AlphaFold3. For the AlphaFold3 baseline mentioned across experiments, we used the official AlphaFold3 weights and source code [2].

To implement the corresponding forward models we used PyTorch (2.3.1+cu121) [26]. However, to efficiently calculate the density for larger structures, we used PyKeOps [6] and PyTorch.% All computations were performed on NVIDIA H100 and L40 GPUs running on Debian GNU/Linux 12 distribution.

Across all experiments, we retrieve the multiple sequence alignments (MSAs) using a wrapper around the ColabFold MMseqs2 API [24]. The wrapper submits the query sequences to the remote MMseqs2 server via HTTP POST, polls for job completion, and downloads and extracts the results. The output is provided in A3M format, which is compatible with AlphaFold3.

### 2.2 Input data

#### 2.2.1 X-ray crystallography

##### Atomic models

In the X-ray crystallography workflow, the input consists of a PDB ID, a chain identifier, and an amino acid subsequence in the chain. For this modality, we restrict our modelling to single protein chains. The corresponding PDB structure and MTZ file is retrieved from PDB-Redo [15] and the specific chain is identified using Gemmi (v0.6.5) [39]. From this chain, we extract only the standard amino acid residues explicitly modeled in the PDB file, while excluding structural waters, hydrogen atoms, and all other non-standard amino acids, and save the resulting coordinates as a separate PDB file. In addition, we mutate Selenium Methionine (`MSE`) to Methionine (`MET`) by replacing Selenium (`SE`) atoms with Sulfur (`SD`). We also mutate S-hydroxycysteine (`CSO`) to Cysteine (`CYS`) by removing the Oxygen from the hydroxyl group (`OG`). If alternate conformations (altlocs) are present [30], the PDB is split into the corresponding altloc files. Since AlphaFold3 uses one-based residue indexing, we renumber the residues in each split PDB to start from 1. In many cases, the retrieved PDB files can contain incomplete or unmodelled residues due to poor quality density. To address this, we use PDBFixer [7] (v1.9.0), which employs standard residue templates from OpenMM [8] force fields, to model the missing atoms in all split PDBs. To avoid steric clashes after imputing the atoms with PDBFixer, we relax the structure using AMBER-14 force field [37] relaxation. The isotropic B-factor for these imputed atoms is set to 100.00. For AlphaFold3, the input sequence is limited to the residues explicitly modelled in the reference chain. To construct the atom mask for applying the substructure conditioner, we use the provided amino acid subsequence to identify residues that will primarily be optimized with the density-based loss versus those optimized using the substructure conditioner.

##### Electron density maps

Our proposed method generalizes across different types of real-space electron density maps. Here, we considered three map types.

- 2*F*_o_ − *F*_c_ **maps:** These maps are rendered using the Phenix [3] `fft` command:

`fft hklin mtz_path mapout map_output_path`

`LABIN F1=FWT PHI=PHWT SIG1=SIGFP VF000 volume f000`

The unit cell volume (`volume`) and the total scattering factor at zero scattering angle *F*(0, 0, 0) (`f000`) are derived from the input atomic model. For a triclinic cell with edge lengths *a, b, c* (in Å) and angles *α, β, γ* (in radians), the unit cell volume is:

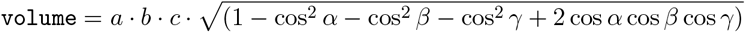

Unit cell parameters (*a, b, c, α, β, γ*) are parsed from the input atomic model. *F*(0, 0, 0) is obtained using `phenix.f000`, which estimates the zero-angle scattering factor by summing electron counts from both the atomic model and the bulk solvent. The bulk solvent contribution is calculated using the default average solvent density of 0.35 e^−^/Å^3^. The atomic and solvent electron contributions are then combined to yield *F*(0, 0, 0).

- **Absolute-scale END maps:** These density maps are generated using the END map workflow of [20], which combines Phenix and CCP4 programs to produce absolute-scale electron density maps (e^−^/Å^3^ units). We run

`phenix.refine atomic_model_file mtz_file strategy=rigid_body`

`/END_RAPID.com pdb_refine_001.eff -norapid`

The atomic model first undergoes rigid-body refinement to prevent artifacts from misaligned models. The END script then rescales the observed structure factors against the calculated ones and offsets the density by adding *F*(0, 0, 0) to every voxel. This places the map on an absolute scale, where zero corresponds to vacuum.

- **Composite omit maps:** These density maps are computed in Phenix using,

`phenix.composite_omit_map atomic_model.pdb reflection_data.mtz`

These maps reduce model bias while minimizing phase degradation.

To align the real-space density maps with the corresponding atomic models from Section 2.2.1, we used `phenix.map_box` with `selection_radius=10` to carve the map around the corresponding atomic model with sufficient padding.

Since our X-ray density guidance operates locally, we carved the input density map to include voxels within a radius of 5Å of all atoms in the residue range (and chain) of interest from the reference PDBs. This pre-processing reduces VRAM memory consumption. During *F*_c_ computation, the ensemble density is calculated at voxels within this 5Å local neighborhood.

#### 2.2.2 NMR

We used three sources of NMR observables for guidance: NOE-implied distance restraints, chemicalshift–implied backbone dihedral angles, and per-residue order parameters.

##### NOE-implied distance restraints

NOE restraint lists were parsed from NMR-STAR files (via `pynmrstar`), selecting only NOE-derived distance restraints; other restraint types such as dihedral angles, H-bonds, and residual dipolar coupling (RDC), were excluded. *Ambiguous NOEs*— cross-peaks whose atom-pair assignment is uncertain—were retained and represented as disjunctive (`OR`) constraints: each ambiguity group defines multiple candidate atom pairs sharing the same bounds, and the restraint is satisfied if *any one* candidate meets those bounds (not necessarily all). Lower/upper bounds were read directly from file; missing lower bounds were set to 0.0 Å. We note that, although NOE physics is intensity-based, we adopt the common semi-quantitative distance-bound representation here, as is standard practice in MD-resolved NMR ensemble workflows; a future extension of our pipeline may incorporate intensity averaging within ambiguity groups prior to distance conversion. For each system, the corresponding NMR-STAR restraint file was retrieved from the PDB entry on the RCSB website.

##### Chemical-shift-implied dihedral angles

Backbone Φ^tgt^/Ψ^tgt^ angles used in combined cryo-EM and NMR guidance were inferred from raw chemical shifts using TALOS-N [32], a tool that infers potential ranges of backbone dihedral angles from chemical shifts. Per target, the chemical-shift lists were retrieved from the BMRB entry cross-referenced to the corresponding PDB ID.

##### Order parameters

Order-parameter (*S*^2^) values were taken on a per-residue basis from the respective publications (or associated supplementary datasets) for each target. Across all observables, when required values or bounds were unavailable for a residue or atom pair, we masked that item and did not guide on it.

#### 2.2.3 Cryo-EM

In the Cryo-EM workflow, the inputs include a reference atomic model, amino acid sequences, and the corresponding electrostatic potential (ESP) map from the EMDB [1].

##### Atomic models

In cryo-EM, the objective is not to refine high-resolution reference structures, as in X-ray crystallography, but to fit amino acid sequences into experimentally resolved volumes. This distinction is because of the lower resolution of cryo-EM ESP maps, which makes them useful for studying larger protein structures. Here, reference PDB structures from RCSB [29] are primarily used to align chain labels, construct atom masks, and serialize outputs. Unlike crystallographic pre-processing pipelines, missing residues or atoms are not imputed with modeling software (like PDBFixer), and alternate conformations are not split into separate models. To ensure consistency with AlphaFold3, all residues are renumbered to be 1-indexed. Finally, because cryo-EM often targets large complexes, multimeric inputs along with multiple sequences can be provided. Relevant residue ranges and chain identifiers are specified alongside the sequences before guidance.

##### Electrostatic potential maps

The input ESP volumes are obtained from the Electron Microscopy Data Bank (EMDB) [1] in ‘.map’ or ‘.mrc’ format. To reduce VRAM usage without losing resolvable information, ESP maps with large side length *D* are downsampled in Fourier space to a smaller side length *D*^*′*^ *< D*. The reported resolution *r* (in Å) is used to verify that the Nyquist frequency of the downsampled grid remains sufficient. This is checked using

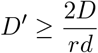

where *d* is the original voxel size in Å. This relation follows from (i) conservation of the total physical side length 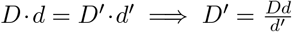 and (ii) the Nyquist criterion 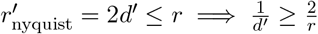.

Lastly, atomic masks are generated using the reference protein to provide controlled guidance, but are constructed in a way that maintains the generality of the approach:

- **Global mask:** A global mask is defined to include all voxels within *r*_max_ = 2.1 Å of the atoms in the reference protein. Empty regions of the masks are filled using convex hull implemented in `scipy.spatial.ConvexHull` [36].
- **Per-chain mask:** For multimeric structures (e.g., amyloids), masks are generated per chain using a similar *r*_max_ distance threshold. Since this is restricted to well-separable chains, userdefined masks could likewise be imported after manual generation. This is later used to obtain better-defined guidance directions.

### 2.3 Force Field

To map experimentally faithful ensembles onto Boltzmann-distributed populations, we evaluate per-structure energies *E*(**X**) using ProteinEBM [28], a learned energy-based model derived from Boltz-2 [25]. ProteinEBM is trained to assign scalar energies to protein conformations at 300 K by learning a Boltzmann-consistent score function over the conformational space. Although ProteinEBM does not constitute a classical force field, it defines an implicit, differentiable energy functional over protein conformations that serves as an effective structural prior for ensemble reweighting.

### 2.4 Protein structure inverse problems

#### Notation

The amino acid sequence of a protein is denoted by **a** and the corresponding 3D Cartesian coordinates of all atoms is denoted by **X** = (**x**_1_, …, **x**_*m*_), where **x**_*i*_ ∈ ℝ^3^ is the position of the *i*-th atom.

#### Problem Setting

Given a protein’s amino acid sequence **a** and its corresponding experimental observation **y** (from crystallography, NMR, or Cryo-EM), our goal is to sample an experimentally faithful ensemble of *n* structures **𝒳**= (**X**^1^, … **X**^*n*^) from the posterior distribution *p*(**𝒳** | **a, y**).

The posterior distribution can be factorized as *p*(**𝒳** | **a, y**) ∝ *p*(**y**, | **𝒳 a**) · *p*(**𝒳** | **a**) using Bayes Rule. The prior term is captured by *p*(**𝒳** | **a**) which is the likelihood of an ensemble conditioned on the amino acid sequence **a**. Using probabilistic laws, individual terms can be factorized into a product of independent samples in the prior term,

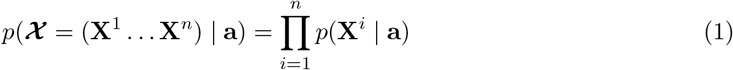

In our case, the prior *p*(**𝒳** | **a**) is implicitly modelled by AlphaFold3 [2].

The data term *p*(**y** | **𝒳, a**) represents the probability of the experimental data **y** given the protein structure ensemble **𝒳** and the amino acid sequence **a**. Since the data term is conditioned on the entire ensemble, we cannot separate the likelihood into separate structures. In essence, the data term models the underlying physics of the experiment. In this paper, we model the data term for three experimental modalities:

1. *X-ray crystallography* where we use real-space electron density (ED) maps;
2. *NMR* where we use nuclear Overhauser effect (NOE) pairwise distance restraints, amide (N-H) and methyl (C-C) order parameter obtained using relaxation (sensitive to ps-ns motion timescale) and residual dipolar coupling (RDC, sensitive to *µ*s-ms motion timescale), and backbone dihedral angles *ϕ* and *ψ* estimated from NMR chemical shifts using TALOS-N [32];
3. *cryoEM* where we use electrostatic potential (ESP) maps.

Additionally, in all modalities we use the following terms:

1. *Substructure conditioning* allowing to fix parts of the structure to follow an existing model;
2. *Validity* ensuring physically plausible bond lengths, valence angles and lack of steric intersections.

In the sequel, we describe in detail the construction of these likelihoods.

#### 2.4.1 Guided diffusion and non-i.i.d. sampling

AlphaFold3 generates structures using a diffusion-based generative model [13] over all-atom coordinates. In its original formulation, a single structure **X** is produced by integrating a variance-preserving stochastic differential equation (SDE) [33].

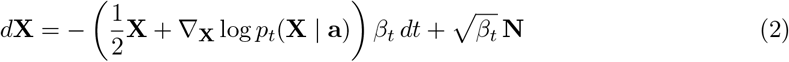

where *β*_*t*_ is the noise schedule, **N** ∼ 𝒩(**0, I**) is sampled from an isotropic normal distribution, and ∇_**X**_ log *p*_*t*_(**X** | **a**) is the learned score function modelled by AlphaFold3 [16]. To predict a single structure, AlphaFold3 samples from the prior distribution **X**_*T*_ ∼ 𝒩(**0**, *β*_0_ · **I**) and integrates the SDE in Equation 2 from *t* = *T* to *t* = 0 to iteratively desnoise the diffusion variable **X**_*T*_ to a noiseless protein structure **X**_0_.

In our setting, we expand the current SDE to model ensembles instead of single structures. Formally, we sample an ensemble **𝒳** = (**X**^1^, **X**^2^, …, **X**^*n*^) of *n* structures. Thus, we can generalize the SDE in Equation 2 to samples ensembles instead of a single structure.

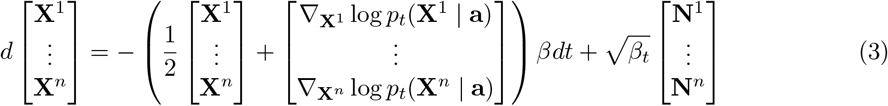

where **N**^*k*^ ∼ 𝒩 (**0, I**) and the unconditional score term 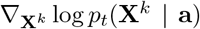 can be separated into independent structures like in Equation 1. In order to sample a *non-I*.*I*.*D*. ensemble from the posterior distribution, we plug in the guidance score to Equation 3, obtaining,

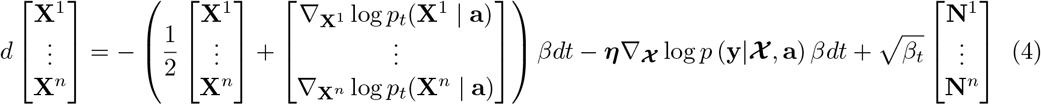

Unlike the unconditional score term, the guidance score term Equation 4 is not separable because it is conditioned on ensemble 𝒳. Also, the hyperparameter ***η*** can be used to scale the strength of the guidance score and direct the flow towards higher posterior likelihood regions. The Pseudocode for guided AlphaFold3 is provided in Algorithm 1.

##### Parameter settings

For X-ray crystallography, we consistently set ***η*** = 0.1. For NOE constraints under NMR, we experimented with values ***η*** ∈ {0.1, 0.2, 0.3, 0.4, 0.5} and selected the best-performing result. For order parameters under NMR, we fixed ***η*** = 0.3. To ensure numerical stability during the diffusion process, we apply gradient clipping to clip the guidance score. For Cryo-EM, the used values were *η* ∈ 0.1, 0.15, 0.2 where the best values per structures were chosen.

### 2.5 X-ray crystallography models

Let 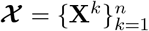 be an ensemble of *n* conformers and conformer **X**^*k*^ have atomic coordinates 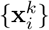 and isotropic B-factors 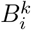. The observed electron density is denoted as *F*_o_ : ℝ^3^ → ℝ. In order to restrict map comparisons around a region of interest, we use a spatial mask.

#### Forward model

To compute the crystallographic likelihood term, we first calculate the theoretical electron density map *F*_c_ : ℝ^3^ → ℝ for all structures in ensemble **𝒳**. *F*_c_ can be calculated at any spatial coordinate location ***ξ*** ∈ ℝ^3^. For a single conformer 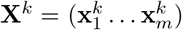 is a sum over kernel-density estimates built from six spherical Gaussians centered at the symmetry operation [5] of every atom.

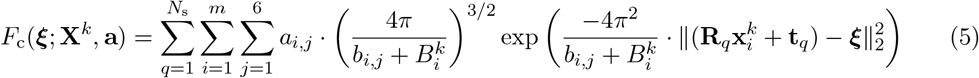

Here, *N*_s_ is the number of symmetry operations [5], *m* is the number of atoms in the asymmetric unit of conformer **X**^*k*^, **R**_*q*_ ∈ *SO*(3) is the rotation matrix of symmetry operation *q*, **t**_*q*_ ∈ ℝ^3^ is the translation vector of the symmetry operation *q, a*_*i,j*_ and *b*_*i,j*_ are tabulated atomic form-factor coefficients for each heavy atom 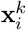 [27], and 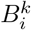 is the B-factor which represents the isotropic displacement parameter [35]. Here, we consider the B-factor to be a bandwidth parameter.

We can extend the density calculation at spatial coordinates ***ξ*** to ensembles

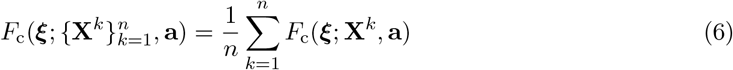

*F*_c_(***ξ***; **X**^*k*^, **a**) is computed using Equation 5. For our experiments, we assume that each atom has a uniform B-factor that is inversely related to the size of the ensemble, 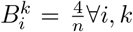. Additionally, we use an ensemble size of *n* = 16 for all our experiments. Our guidance framework differentiably incorporates all symmetry mates of the non-hydrogen atoms belonging to the chain of interest. In subsequent post-processing steps, we further append structural waters, non-standard residues, ligands, ions, and atoms from other chains of the proteins together with their respective symmetry mates.

#### X-ray log likelihood

To quantify the agreement between observed and calculated electron density maps, we measure the *L*_1_ norm of the difference between the calculated ensemble-averaged density *F*_c_ (from Equation 6) and the observed density *F*_o_:

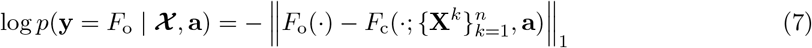

The log-likelihood computation has been summarized in Algorithm 2.

### 2.6 NMR models

#### Differentiable hydrogen placement

AlphaFold3 does not explicitly model hydrogen atoms, yet their positions are essential for accurately modeling NMR observables. To address this, at each sampling step we first reconstruct hydrogen atoms of each atom in an amino acid in a differentiable manner using a PyTorch port of Hydride’s [19] hydrogen placement algorithm. Each hydrogen-bearing center is identified by its local bonding environment and assigned a matching reference fragment. The fragment’s non-hydrogen atoms are then aligned to the target by a single rigid-body superposition, after which the same transformation is applied to its hydrogens. In this way, proton positions adjust smoothly with the heavy-atom coordinates, yielding accurate and fully differentiable NMR calculations.

#### NOE distance restraints

We used solution-state NOE restraints as ensemble-averaged distance constraints. Conceptually, NOEs arise from through-space dipolar interactions between protons with observable effects typically for inter-nuclear separations < ∼ 6 Å; intensities in NOESY spectra reflect an (approximate) *r*^−6^ dependence and are commonly used semi-quantitatively to bound distances. For an ensemble 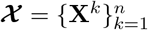, we modeled the ensemble-average distance:

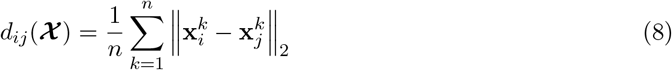

and enforced lower/upper bounds [ine q]via the log-likelihood:

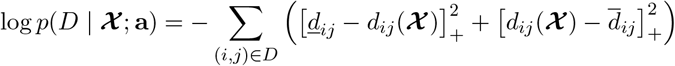

where [*x*]_+_ = max(*x*, 0). Additionally, ambiguous NOE assignments were retained as OR groups; for evaluation, the group-level violation equals the minimum violation among its members (i.e., the group is satisfied if any member lies within bounds). For restraints involving methyl groups, we modeled rapid threefold internal rotation by averaging over the three methyl proton positions generated by differentiable H placement.

We guided the diffusion sampler with the NOE likelihood using non-i.i.d. ensemble sampling, which couples conformers through the ensemble-level likelihood. Substructure conditioning was not used for NOE.

#### Order-parameters

Order parameters *S*^2^ for amide and methyl bonds represent ensemble flexibility (lower values imply higher disorder). At each sampling step, we rigidly align all conformers **X**^*k*^ ∈ 𝒳 in the ensemble to a reference **X**^ref^ to remove global rotation and translation. For each bond *p* in ensemble member **X**^*k*^, we compute a unit bond vector 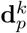(amide uses the N→H vector while methyl uses the C→C vector pointing to a methyl carbon; some amino acids like leucine, isoleucine and valine have two methyl groups and, hence, two vectors per residue). Given these normalized vectors, the calculated bond-order 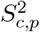 of the bond *p* is given by:

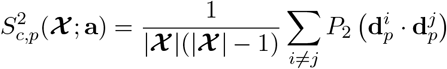

where 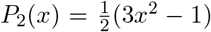 is the Legendre polynomial of order 2. The log-likelihood is computed according to

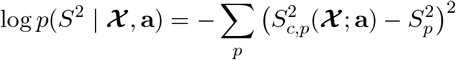

where the summations is performed over the collection of bonds on which the experimental data 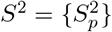 have been collected.

#### Dihedrals likelihood

Backbone dihedral restraints assess the agreement between predicted Ramachandran angles (*ϕ*^pred^, *ψ*^pred^) from ensemble ***X*** and target angles (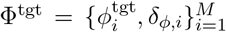 and 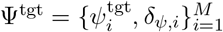) defined over *M* residues. The target angles and their associated uncertainty half-widths are obtained from TALOS-N [32] predictions. The log-likelihood is computed using a 2*π*-periodic (wrapped) angular discrepancy function that respects the circular nature of angles. This term encourages locally accurate backbone geometry and regular secondary structure, complementing global guidance terms derived using NOE distance restraints or Cryo-EM ESP maps. The dihedral log-likelihood is defined as:

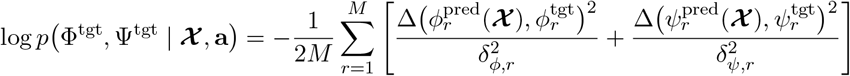

Where, Δ(*α, β*) denotes the wrapped angular difference (implemented as `angle_diff`). This log-likelihood is averaged over all chains of structures in the ensemble.

#### Overall NMR likelihood

When guiding with order-parameters, NOE observations, and dihedrals, the log-likelihoods are combined as

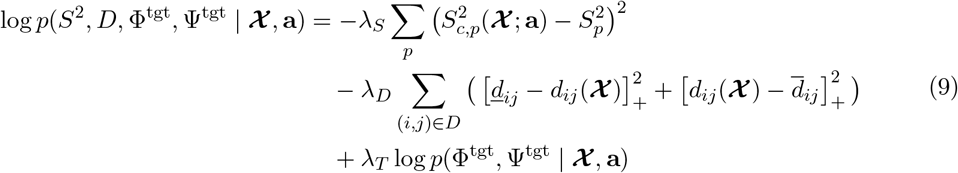

We guide the model either without order parameter information, or using one of two types: relaxation-derived order parameters (ps-ns motions) or residual dipolar couplings (RDC)-derived order parameters (*µ*s-ms motions). Dihedral restraints are incorporated only in experiments that involve Cryo-EM ESP maps. The scaling parameters (*λ*_*S*_, *λ*_*D*_, *λ*_*T*_) are set as follows:

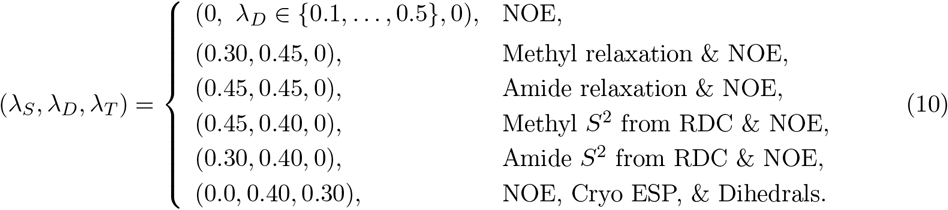

When guiding with NOE’s alone, we selected the best-performing value of *λ*_*D*_ from the tested set.

### 2.7 CryoEM models

#### ESP forward model

As discussed in [38], the forward model for cryoEM electrostatic potential (ESP) maps is approximated by atomic scattering potentials, following equations (5) and (6). Unlike X-ray crystallography, cryoEM maps typically represent a single particle, so the model reduces to

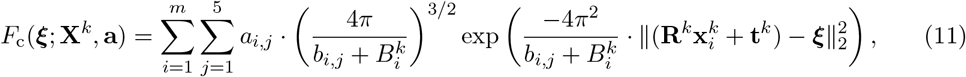

where *ϕ* = (**R**^*k*^, **t**^*k*^) represents the pose of conformation **X**^*k*^ instead of a crystallographic symmetry operator. In the presence of significant blurring, the ensemble contribution can be evaluated as an average using Equation 6.

#### Protein – ESP map alignment strategies

AlphaFold3’s noisy time marginals **X**_*t*_ (atom coordinates at timestep *t* of the diffusion process) and the denoised predictions 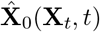 evolve on the learned diffusion manifold and not in a fixed frame. As a result, accurately predicted structures will generally be misaligned with the ERP map’s voxel grid. Additionally, the 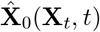 predictions have a random pose at every iteration. In X-ray crystallography, Kabsch Algorithm [17] is used to solve the alignment problem. On the other hand, in cryo-EM *ab initio* protein-density fitting, no reference protein structure is available, making this alignment method inapplicable. Instead, a pure protein-density alignment strategy has to be implemented. Depending on whether the diffusion is performed over the whole ESP map or over well-separable chains, two different alignment strategies have been implemented:

- **Protein – full ESP map rigid alignment**: Global pose determined by solving:

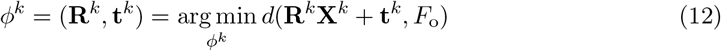

where *d*(**X**, *F*_o_) is the error objective defined in the following sections. The resulting pose is applied as a rigid transformation, 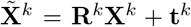 and all subsequent guidance scores are evaluated with respect to the aligned structure.
- **Protein – per chain non-rigid ESP alignment**: In this case, the alignment is performed independently for each chain. A set of per-chain poses 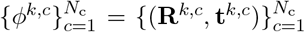 is estimated and applied to obtain the aligned structure 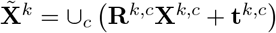, where *N*_c_ is the number of chains.

In AlphaFold3, a structure **X**_*t*_ is represented as a tensor of shape (*N*_a_, 3) where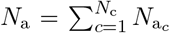, denotes the total number of atoms and 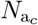 denotes the number of atoms in a chain *c*. After concatenation of 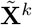, the application of the per-chain poses is a non-rigid transformation with respect to the full structure **X**^*k*^.

The chosen objective *d* is the trilinear interpolation 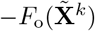, and the maximization of *F*_o_ was implemented as a gradient descent over the pose *ϕ*^*k*^. For this, hyperparameters of number of iterations *N*_iter_, *N*_init_ initial random rotation, learning rates *α*_R_, *α*_*t*_, the side length *T* of the box around the weighted centroid of *F*_o_ and the positive B-factor *B* to be applied to the volume to ensure non-zero gradients for initial poor alignments can be defined in the configuration file. For cases where predictions 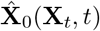 have a starkly different tertiary structure from the intended **X**^*k*^ and this creates numerous unstable local minima (e.g., like what we observed in the insulin receptor structures `8U4B` and `8U4E`, a momentum decay *α*_*m*_ can be specified. This adds decaying Kabsch-aligned anchoring to the pose from the previous diffusion iteration to prevent large fluctuations of poses between diffusion time steps.

#### After-alignment diffusion manifold correction of X_*t*_ **and** 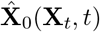

Since in the diffusion process, **X**_*t*−1_ is computed as a function of the noisy structure and the *t* = 0 prediction 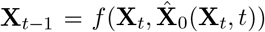, in case a transformation was applied to the aligned 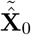, for the correct **X**_*t*−1_ evaluation, an equivalent change must be applied to **X**_*t*_ in its relative frame.

- **Rigid alignment correction:** here the correction can be avoided if the direction gradient is evaluated using autograd as 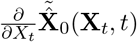, and the aligned 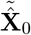 is aligned back through 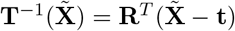.
- **Non-rigid alignment correction:** For correction and preservation of the manifold, non-rigid alignment is applied in two steps: (1) a rigid alignment **T**_1_(**X**) = **R**_1_**X** + **t**_1_ that aligns the complete **X** with the complete density *F*_o_, and (2) the non-rigid per chain *c* transformations 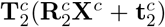 computed using gradient ascent. In this way, movements equivalent to the additional non-rigid fine-detail alignments **T**_2_ of 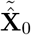 can be defined in the relative frame of **X**_*t*_ and 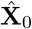as 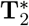, where the corrected structures we will denote as 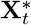 and 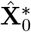. In other words, 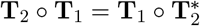. Thus, 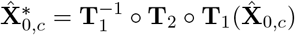 and 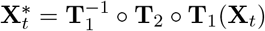

#### ESP *L*_1_ likelihood

The objective *L*_1_ is defined similarly to equation (7) and is computed as −∥*F*_c_(**X**) − *F*_o_∥_1_. This likelihood is best suited for homogeneous proteins or for heterogeneous ones with modest flexibility.

#### Optimal transport likelihood

Different cryoEM ESP maps for a single sequence or protein assembly typically represent distinct states of a heterogeneous protein. Depending on the range of movement between the conformations of such proteins and the original predictions of AlphaFold3, the *L*_1_ density objective gradients are often unable to guide the AlphaFold3 prediction to the map. Instead, due to floating point limitations and the absence of gradients for corresponding atoms that are far apart, this guidance strategy tends to push the noisy structures **X**_*t*_ and the corresponding 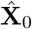 predictions off the diffusion manifold. Therefore, a better approach for evaluating guidance gradients is to use a likelihood that directly points towards the density mass — namely, optimal transport.

We model the protein and ESP map as discrete measures 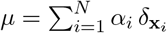 and 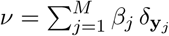, where **x**_*i*_ ∈ ℝ^3^ are all the atoms of a protein and **y**_*j*_ ∈ ℝ^3^ are all the voxel centers inside the global mask (or equivalently, per-chain masks and per-chain coordinates). *α*_*i*_ is proportional to the atomic number *Z*_*i*_ and *β*_*j*_ is proportional to the observed value in the center of the voxel of *F*_0_. The total mass of ***α*** and ***β*** is normalized as 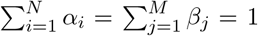. The ground cost matrix is defined as **C** ∈ ℝ^*N ×M*^ with *C*_*ij*_ = *c*(**x**_*i*_, **y**_*j*_) proportional to the 1- or 2-Norm between these two vectors. The *entropy optimal transport* objective is then defined as the *primal problem*:

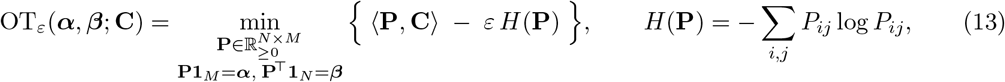

with ⟨**P, C**⟩ = ∑_*i,j*_ *P*_*ij*_*C*_*ij*_ and **1**_*K*_ the *K*-vector of ones. Here **P** is a transport plan, where *P*_*ij*_ specifies the proportion of mass from source atom **x**_*i*_ assigned to voxel **y**_*j*_. The entropy term *εH*(**P**) promotes smooth, stable couplings and well-behaved gradients via discretizing assignments that are too sparse or sharp (coctain a lot of zero entries) and makes the problem strictly convex.

In practice, to reduce computational cost, we use the *Sinkhorn entropy optimal transport loss* in its dual representation with atom/voxel potentials ***f*** ∈ ℝ^*N*^ and ***g*** ∈ ℝ^*M*^ :

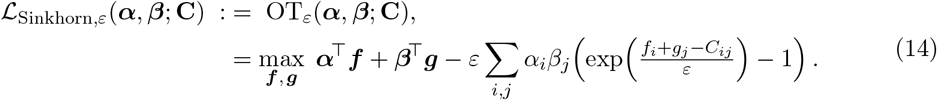

We chose a powerful optimized implementation of optimal transport from `geomloss` [10], particularly to avoid memory bottlenecks. The hyperparameters of choice were `p=2`, `blur=ϵ=0.005`, and the binary `reach` flag, depending on whether mass needed to be dropped for a structure.

In cases where ESP maps contain separable chains, the OT objective was also applied per chain. The rationale was to separate the guidance directions and to prevent unwanted misaligned stable local minima. This was required for structures `6W0O`, `7DA4`, and `9FH1`. A typical issue with those structures was the discrepancy in AlphaFold3’s tensor representation and the spatial arrangement of the subunits, where those were not ordered correspondingly. To solve that, we tested both the Hungarian algorithm for the Linear Assignment Problem (LAP), defined by chainwise distances and brute-force approaches. However, the most optimal solution was the per-chain OT objective: such guidance naturally ordered the AlphaFold’3 tensor representation to the spatial chains and thereby optimally solved for the correspondence implicitly.

#### Recycling and extended optimization of proteins

In cases where AlphaFold3 early-time or unguided predictions poorly match the protein of interest, the initial protein optimization required to reach sufficient quality for ESP map alignment may consume the majority of available time steps in the diffusion process. Such a trend was observed for structures `6W0O` and `9FH1`. This, in turn, can leave an insufficient number of steps in the diffusion process for fine-detail protein-ESP map fitting, or make it completely impossible due to the noise schedule being insensitive to any change after the noise level *σ*_*t*_ reaches its lowest values (in practice, no significant change is observed after 160 steps). Additionally, the aforementioned proteins suffer from strong, stable local minima in the OT likelihood, e.g., 180^°^ rotations about any axis lying on the plane perpendicular to the first principal component of `6W0O`. This significantly increases the average time it takes for the protein to align to the intended spot during the diffusion process.

To allow additional time for finding these optimal protein-ESP map arrangements, we implement recycling: once the noisy structure reaches a specified time step *t*, instead of proceeding with the backwards diffusion process to obtain 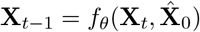, noise is added as in the forward diffusion process to advance from time step *t* to *t* + *k*:

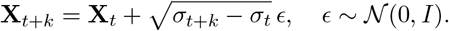

For example, if protein-ESP alignment of `6W0O`converged to all the intended rotations for each chain instead of the 180^°^ opposite-symmetry poses after 160 time steps, it would not be further optimized. Adding an additional small level of noise, typically equivalent to 25 time steps of the forward process, does not break the correctly found tertiary structure while giving extra time for fine-detail fitting.

#### Objective normalization

To balance multiple objectives with the ESP guidance, we employ a specific normalization strategy: one objective, typically the ESP *L*_1_ or OT likelihood, is chosen as the reference, and all other likelihood terms are rescaled at each iteration so that their contributions remain a fixed proportion *λ*_*i*_ ∈ [0, 1] of this main objective. This prevents magnitude imbalances that are otherwise observed when combining losses from very different modalities, and ensures stable relative weighting throughout the diffusion process.

#### Overall Cryo-EM likelihood

The overall (pure) Cryo-EM guidance objective is defined as a weighted combination of the density-based *L*_1_ and the OT likelihood:

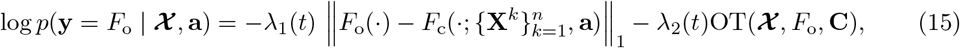

where *λ*_1_(*t*) and *λ*_2_(*t*) are time-dependent weights normalized as described above: one of these weights is set to *λ*_*i*_ = 1 as the main objective while the other is some portion of it normalized as per previous section. For the pure Cryo-EM loss, we typically simply use *λ*_1_ = 0 and *λ*_2_ = 1 until the structure roughly aligns to the density mass and then switch to pure density loss to optimize fine-detail features and optimize B-factors, i.e., *λ*_1_ = 1 and *λ*_2_ = 0. Some more complicated mixtures were explored; however, they showed better results for combined guidance with other modalities. For example, in the NMR setting, the likelihood is scaled to 0.15 of the active objective, first relative to the OT loss (when it is turned on), and later relative to the *L*_1_ density loss (once it becomes active).

### 2.8 Additional data terms

#### Substructure conditioner

Specifically for crystallographic structures, we aim to optimize a certain region of the protein while the remainder of the structure is stabilized by bootstrapping the diffusion process to a set of reference atomic coordinates. This is similar to `SubstructureConditioner` in Chroma [14].

As input, consider a list of reference atom locations *Y* = {**y**_*i*_ : *i* ∈ *A*} for atom indices *A* ⊆ {1 … *m*} where *m* is the number of atoms in conformer **X**^*k*^. The log-likelihood is as a quadratic penalty on the deviation from reference atom locations

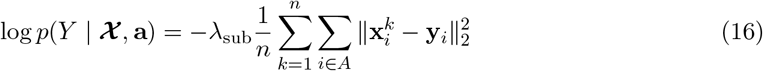

Before computing the log-likelihood we align all structures in the ensemble to the reference atom locations (limited to indices in *A*) using Kabsch algorithm [17]. For crystallography, *λ*_sub_ = 0.1, while for NMR and Cryo-EM, *λ*_sub_ = 0.0.

#### Validity likelihood

In order to prevent ensembles with elongated bonds and steric clashes, we incorporate a validity log-likelihood as regularization akin to AlphaFold2’s violation loss [16]. Consider an ensemble of *n* protein conformations 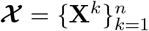, where each structure **X**^*k*^ ∈ ℝ^*m*×3^. We define a binary bond matrix **B** ∈ {0, 1}^*m*×*m*^ such that *B*_*ij*_ = 1 if atoms *i* and *j* are covalently bonded, and 0 otherwise. The bond length loss for conformation **X**^*k*^ over bonded atom pairs (*B*_*ij*_ = 1) is given as,

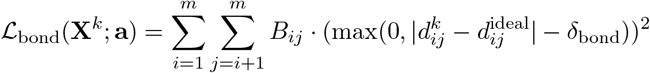

where, 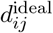 is the ideal bond length approximated as the sum of covalent radii of atoms *i* and *j*, 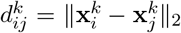 is the between atoms *i* and *j* in conformation *k*, and *δ*_bond_ = 0.2Å is a tolerance margin. For non-bonded pairs (between and within residues), steric clashes are penalized using a soft collision loss, defined as the maximum violation over neighbors of each atom,

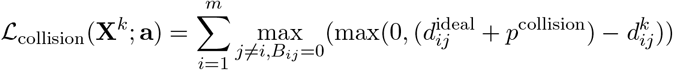

Here, we use a padding distance *p*^collision^ = 0.4 (in Å) to prevent over-penalization of near-contact atoms. Lastly, we define a bond-angle violation loss as

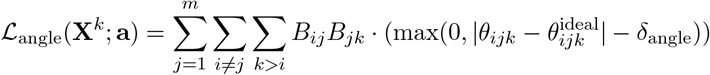

Where *θ*_*ijk*_ (in degrees) is calculated using the dot product of the bond vectors from central atom *j* to atoms *i* and *k*, 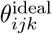 (in degrees) is retrieved from the Valence Shell Electron Pair Repulsion (VSEPR) theory [12], and *δ*_angle_ = 12^°^ is a tolerance margin. The resulting validity log-likelihood across the ensemble is given as

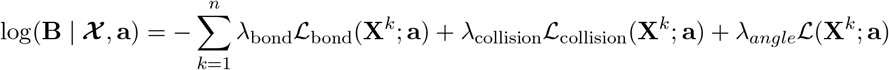

Where, *λ*_bond_, *λ*_collision_, and *λ*_angle_ are scaling factors that control the contribution of bond, collision, and bond angle terms. For ensembles guided using crystallographic density maps, we use *λ*_bond_ = *λ*_collision_ = *λ*_angle_ = 0.075. For NMR-guided ensembles, the scaling factors (*λ*_bond_, *λ*_collision_, *λ*_angle_) depend on the choice of (*λ*_*S*_, *λ*_*D*_, *λ*_*T*_): ^1^

### 2.9 Force-field relaxation

Despite applying validity likelihood, a select few samples have broken bonds or steric clashes. Due to the non-physical nature of the ensemble, we filter out these samples from the ensemble. We consider a sample to have broken bonds if the distance between any pair of bonded atoms exceeds *τ*_bond_. Likewise, we consider a sample to have steric clashes if the distance between any two atoms is less than *τ*_clash_. Across all experiments, *τ*_bond_ = 2.1Å and *τ*_clash_ = 1.1Å.

Despite this, certain geometries can go undetected. To resolve this while maximizing the log-likelihood of the experimental observations, we relax the remaining samples in the ensemble using an off-the-shelf harmonic force-field. Specifically, we use OpenMM’s [8] implementation of the AMBER14 [37] force field parameters for energy minimization within ColabFold [24]. In ColabFold, the energy is minimized for a maximum of 2000 iterations with energy tolerance threshold of 2.39 kcal/mol and stiffness of 100.0 kcal / mol Å^2^ for NMR and 10.0 kcal / mol Å^2^ for the other modalities, thereby controlling the strength of positional restraints applied to the atoms.

### 2.10 Ensemble pruning

#### Note

Here, *ensemble* refers to the set of samples that remain after relaxation, from which we prune a non-redundant subset. After relaxation, we aim to report a non-redundant subset of samples ***X*** _ℐ_ = {**X**^*k*^ : *k* ∈ ℐ} that best explains experimental observation **y**. ℐ denotes the indices of selected samples from the full ensemble. We propose two complementary strategies for populating ℐ.

#### Orthogonal matching pursuit (OMP)

OMP [23] is a greedy selection algorithm that begins with ℐ = ∅. At each iteration, the candidate *k* ∉ ℐ that maximizes log *p*(**y** | **𝒳** _ℐ∪{*k*}_, **a**) is added to the support ℐ set until the log likelihood no longer increases. After termination, each sample in **𝒳**ℐ is assigned a uniform occupancy of 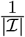.

To avoid overfitting to noise in **y** and reduce redundancy in the ensemble, we introduce two early-stopping criteria. (i) The ensemble size is capped at *n*_max_ = 5. Once |ℐ|= 5, the algorithm terminates. (ii) Before including a candidate, we evaluate the ensemble’s validation loss *f*_val_, and the algorithm terminates if the loss increases. The OMP selection procedure is applied only to ensembles derived from X-ray crystallography.

The selection of the best-performing sample at each iteration is based on Equation 6. The log-likelihood is computed over all voxels ***ξ*** in *F*_o_ that are within 2.5 Å of atoms in the relevant residue range of the reference PDB files (Section 2.2.1). Before a candidate is added to **𝒳**_ℐ_, a scalar B-factor *B* (applied uniformly to all atoms in the structure) is optimized to maximize the log-likelihood. This effectively tunes the bandwidth of *F*_c_(***ξ***; **𝒳**_ℐ_, **a**) to best match the observed density *F*_o_(***ξ***). Optimization of *B* is carried out using Adam [18] for 100 iterations with a learning rate of 1.0. As a validation loss, we use *R*_free_ (Section 2.11), computed with Phenix’s `phenix.model_vs_data`.

#### Occupancy optimization (OCC)

*In crystallography*, we propose an alternative pruning strategy since OMP does not always yield the minimal subset explaining *F*_o_. Instead, we optimize conformer occupancies in the relaxed ensemble **𝒳**_ℐ_ to best fit *F*_o_. Each candidate initially assigned a uniform weight 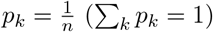 with *n* samples in the ensemble **𝒳** = (**X**^1^ … **X**^*n*^).

In this algorithm, given **𝒳** and amino acid sequence **a**, the *weighted average* calculated electron density is defined as

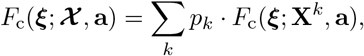

where *F*_c_(***ξ***; **X**^*k*^, **a**) is the calculated density of conformer **X**^*k*^ weighted by occupancy *p*_*k*_. The occupancy weights are optimized by minimizing the following objective:

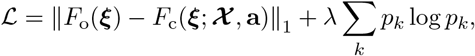

The first term maximizes the log likelihood log *p*(*F*_o_ | **𝒳, a**), while the second term serves as a sparsity regularizer that promote a small subset of conformer retain significant occupancy. The objective is minimized using Adam optimizer for *T*_max_ = 300 iterations with a learning rate of *α* = 10^−3^ and *λ* = 0.7. We observe that, most occupancies decay to zero thereby best explaining *F*_o_ with minimal redundancy. After convergence, the pruned ensemble is defined as

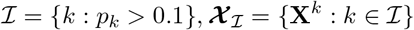

A detailed pseudocode is provided in Algorithm 6. In practice, we apply both algorithms and ultimately select the minimal ensemble that provides the best fit to *F*_o_.

#### Refinement

*In crystallography*, after the termination of the ensemble pruning algorithms, the selected ensemble **𝒳** _ℐ_ is merged into a single PDB file. In this merged structure, atoms included in the set *A* (outside residue range of interest) are retained from the corresponding chain of the PDB-Redo entry [7], while atoms not included in *A* (in residue range of interest) are replaced with those modeled in **𝒳** _ℐ_. Each conformer in **𝒳** _ℐ_ is assigned a unique alternate conformer (altloc) identifier (*A, B*, etc.). Altloc occupancies are then set according to the probabilities estimated by the selection algorithms (see Algorithms 6, 5). Other structural attributes, such as isotropic and anisotropic B-factors, are retained from the relaxed conformers.

The merged structure is then refined against the experimental diffraction data using `REFMAC5`, part of the CCP4 library [4]. `REFMAC5` refines a structural model by minimizing the difference between the observed structure factors (provided in the MTZ file from PDB-Redo) and the calculated structure factors (computed from the input model). The refinement program adjusts atomic coordinates and atomic displacement parameters (B-factors) to better fit the observed structure factors. During optimization, `REFMAC5` applies stereochemical restraints to maintain chemically plausible geometry. Since the coordinate updates are typically small, the refinement step preserves the conformational diversity introduced by our guidance framework while ensuring statistical consistency with the diffraction data. In practice, we perform 5 cycles.

### 2.11 Metrics

#### 2.11.1 X-ray crystallography

To evaluate how well our density-guided structures agree with experimental X-ray crystallographic data, we report three metrics: R values, RSCC, and cosine similarity.

##### R-factors

The R-factor (or crystallographic R-value) is a standard metric for assessing how well a crystal structure’s model explains experimental X-ray diffraction data. This data is typically stored in MTZ files. The metric measures the alignment between the observed structure factor amplitudes |*F*_obs_| and the calculated structure factor amplitudes |*F*_calc_|. Formally,

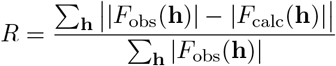

Here, **h** denotes a Miller index triple (*h, k, l*) which denotes a discrete reflection in the reciprocal (Fourier) space. To avoid overfitting, *R*_free_ value is calculated in a similar manner, but using a small subset *T* of reflections that are excluded from the refinement pipeline and consequently serve as a validation set. Hence,

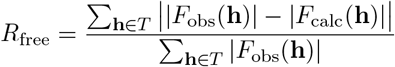

It is important to note that |*F*_obs_| and |*F*_calc_| are structure factor amplitudes in reciprocal space. On the other hand, *F*_o_ and *F*_c_ used in our density-guidance framework denote the real space 3*D* volumetric electron density grids. Electron density maps are obtained by estimating phase to the structure factor amplitudes followed by an inverse Fourier transform into real space. For each experiment, we report both the R and *R*_free_ values as calculated by `REFMAC5` after refinement.

##### Real Space Correlation Coefficient (RSCC)

The real-space correlation coefficient is a local validation metric that measures the alignment between a structural model and the observed experimental electron density map. Unlike R-values, RSCC is calculated directly in real space by comparing observed electron density *F*_o_(***ξ***) with calculated electron density *F*_c_(***ξ***) over an amino acid. Formally, RSCC for a grid of voxels 𝒢 around a residue *a* ∈ **a** is given by,

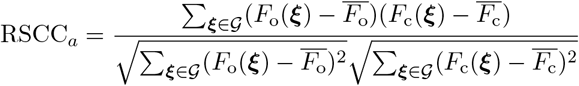

Where, 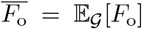, and 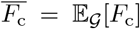. RSCC ranges from −1 to 1, with values closer to 1 indicating a residue that is well supported by the experimental density. For our use case, we used `phenix.real_space_correlation` to measure the RSCC for each residue (and altloc) in our protein.

##### Cosine Similarity

Whereas RSCC provides a residue-level measure of model-to-map alignment in real space, we are also report in a single global score over the entire residue range of interest. For this purpose, we use cosine similarity to quantify the overall alignment between the observed density *F*_o_ and the calculated density *F*_c_. To compute this metric, we first identify the set of spatial coordinates ***ξ*** ∈ ℝ^3^ by selecting all voxels from *F*_o_ that lie within 2.5Å of atoms in the relevant residue range of the reference PDB structures. Formally,

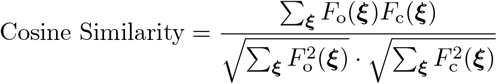

where *F*_o_(***ξ***) and *F*_c_(***ξ***) denote vectors of density values at the selected voxel coordinates. As with RSCC, values closer to 1 indicate strong agreement between calculated and observed densities, signaling a good overall structural fit to the experimental map.

#### 2.11.2 NMR

##### Percentage of violated NOE constraints

Given a set of NOE distance-bound constraints grouped into ambiguous OR-groups, 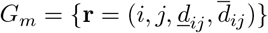, define the ensemble-averaged interproton distance

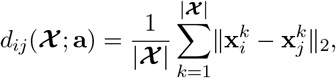

and the per-restraint violation magnitude

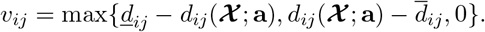

The group-level violation is 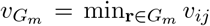, so an OR-group is satisfied if any member falls within bounds. The percentage of violated NOE constraints is

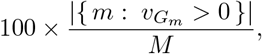

where *M* is the number of OR-groups. Ambiguity and methyl restraints are treated as in the NOE likelihood (ensemble-averaged distances; methyls averaged over the three proton positions).

##### Median violation magnitude

Given the group-level violations 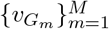 defined above, the median violation magnitude is

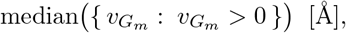

i.e., the median taken over violated groups only.

##### Order parameter correlation (*r*-factor)

Given a set of experimentally determined order parameters, 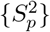 for some collection of bonds *p* ∈ *P*, and a corresponding set of order parameters 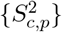 estimated from the generated ensemble, the correlation coefficient is defined as

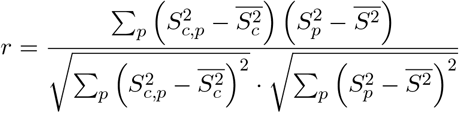

where 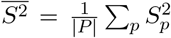 and 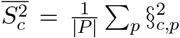 are the averages of the experimental and estimated order parameters, respectively. The coefficient ranges from −1 to 1, with higher values indicating better agreement with the experiment.

##### Order parameter normalized fitting error (*q*-factor)

Similary to the correlation coefficient, the normalized fitting error is defined as

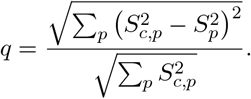

Lower values of *q* indicate a better agreement with the experiment. While *r* is insensitive to a constant offset in the estimated values, *q* is sensitive both to the error standard deviation and bias.

##### ANSURR metrics

ANSURR [11] (Accuracy of NMR Structures Using RCI and Rigidity) validates a protein model by comparing per-residue flexibility inferred from backbone chemical shifts with rigidity predicted from the 3D structure. Flexibility from NMR data is estimated via the Random Coil Index (RCI), while structural rigidity is computed from the network of covalent and hydrogen-bond constraints using rigidity theory (e.g., FIRST). ANSURR reports two centile-ranked scores (0-100): a *correlation* score that reflects agreement in the pattern of flexible vs. rigid regions, and an *RMSD* score that reflects agreement in the magnitude of rigidity/flexibility. For structural ensembles, ANSURR is computed separately for each ensemble member, and the median of RMSD and correlation scores across the ensemble are used as representative metrics.

#### 2.11.3 Cryo-EM

For Cryo-EM, the same metrics as for Crystallography were taken, mainly the cosine similarity and the per-residue model-map cross-correlation (CC) scores from Phenix from section 2.11.1.

### 2.12 Minimal end-to-end code invocation

#### 2.12.1 X-ray crystallography

To guide AlphaFold3 predictions using real-space crystallographic electron density maps, run the following command in your terminal:

`python3 run_xray.py <pdb_id> <chain_id> <region_subseq> \`

`--ccp4_setup_sh <CCP4 SETUP PATH> --phenix_setup_sh <PHENIX SETUP PATH>`

Here, `<pdb_id>` is the four-character PDB identifier of the protein, `<chain_id>` specifies the target chain, `<region_subseq>` is the subsequence of the amino acid sequence to be guided. The `--ccp4_setup_sh` and `--phenix_setup_sh` arguments point to the respective CCP4 and PHENIX installation scripts.

#### 2.12.2 NMR

To guide AlphaFold3 predictions using NOE distance restraints, run the following command in your terminal:

`python3 run_nmr.py <pdb_id>`

Here, `<pdb_id>` is the four-character PDB identifier of the protein. Optionally, users may also supply methyl RDC and relaxation files (`--methyl_relax_file` and `--methyl_relax_file`), as well as amide RDC and relaxation files (`--amide_rdc_file` and `--amide_relax_file`), to provide additional experimental guidance during structure prediction.

#### 2.12.3 CryoEM

To guide AlphaFold3 predictions using CryoEM electrostatic potential maps, run the following command in your terminal:

`python3 run_em.py <pdb_id> <emdb_id> <renumbered_pdb_file> <assembly_identifier> \`

--phenix_setup_sh <PHENIX SETUP PATH> --dihedrals_file <TALOS dihedrals> \

--noe_restraints_file <NOE restraints> --noe_pdb_file <renumbered NOE file path> \

--sequences <space separated sequence> --counts <#chains for each sequence>

Here, `<pdb_id>` is the four-character PDB identifier of the protein, `<emdb_id>` is the EMDB [1] identifier of the cryo-EM electrostatic potential map and `<renumbered_pdb_file>` is the path to a PDB file whose residues have been renumbered to match the 1-indexed amino acid sequences. In other words, if residue A in the input sequence is at index 5, the corresponding residue in the PDB must also be renumbered to index 5. Missing or unresolved atoms are permitted. Since the modeling is performing *ab-initio*, the PDB is not used during guidance. Instead, it is used exclusively for evaluation and for constructing the initial volume mask when the default density when the default density-based alignment is selected (as opposed to RMSD alignment to atomic coordinates). Multichain compositions are specified with `--sequences` (space-separated amino-acid sequences for each unique chain) and `--counts` (the corresponding copy numbers, in order). `<assembly_identifier>` specifies the biological assembly or asymmetric unit to model. The `-phenix_setup_sh` argument point to the PHENIX installation script. Our framework also supports joint guidance using ESP maps. These optional restraints can be provided via `--dihedrals_file` (e.g., TALOS-N dihedrals), `--noe_restraints_file` (NOE distance restraints), and `--noe_pdb_file` – in the similar fashion to the renumbered pdb file, all the residues in these files should be reordered to match the 1-indices in the sequences given to our guidance model.

### 2.13 Energy-reweighted Guidance

While the ensembles generated using the above method are experimentally faithful, they do not reflect equilibrium population weights implied by an underlying physical energy model. For this, we introduce an additional reweighting step that maps experimentally faithful ensembles onto approximately Boltzmann-distributed populations. We assume access to an energy function *E*(**X**), derived from a force-field model which induces a Boltzmann density.

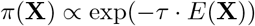

Where *τ* = (*k*_*B*_*T*)^−1^, *k*_*B*_ = 0.001987 kcal mol^−1^K^−1^ is the Boltzmann constant, and *T* is the temperature in Kelvin (*K*). Using this Boltzmann density, we modify the ensemble statistic in 8 to define an energy-weighted ensemble statistic 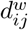,

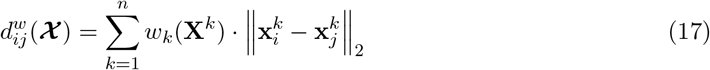

Where the normalized Boltzmann weights are given by 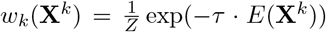 with 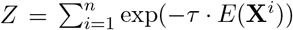 as the normalization constant. Substituting this weighted ensemble statistic into the NMR log-likelihood yields

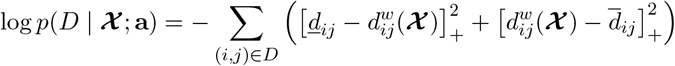

Guiding with this log-likelihood biases the ensemble toward structures that are both experimentally consistent and energetically favorable.

To balance exploration and energetic refinement during the diffusion process, we anneal *τ* over time *t* according to

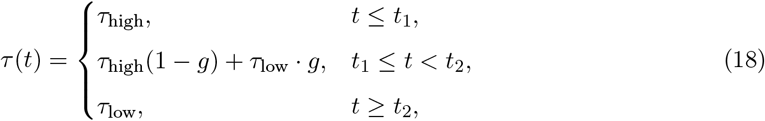

where

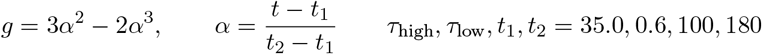

We set *τ*_high_ = 35, *τ*_low_ = 0.6, *t*_1_ = 100, *t*_2_ = 180. This schedule enables early-stage exploration under a flattened energy landscape, followed by progressive sharpening toward low-energy configurations.

Finally, to mitigate noise and short-range artifacts in the force-field energy estimates during late-stage refinement, we apply an exponential moving average (EMA) to the energies after 160 diffusion steps:

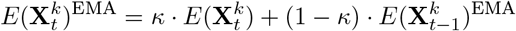

With smoothing parameter *κ* = 0.3. This improves the robustness of energy-based guidance without suppressing structural diversity.

#### Algorithm 1

AlphaFold3 Guidance

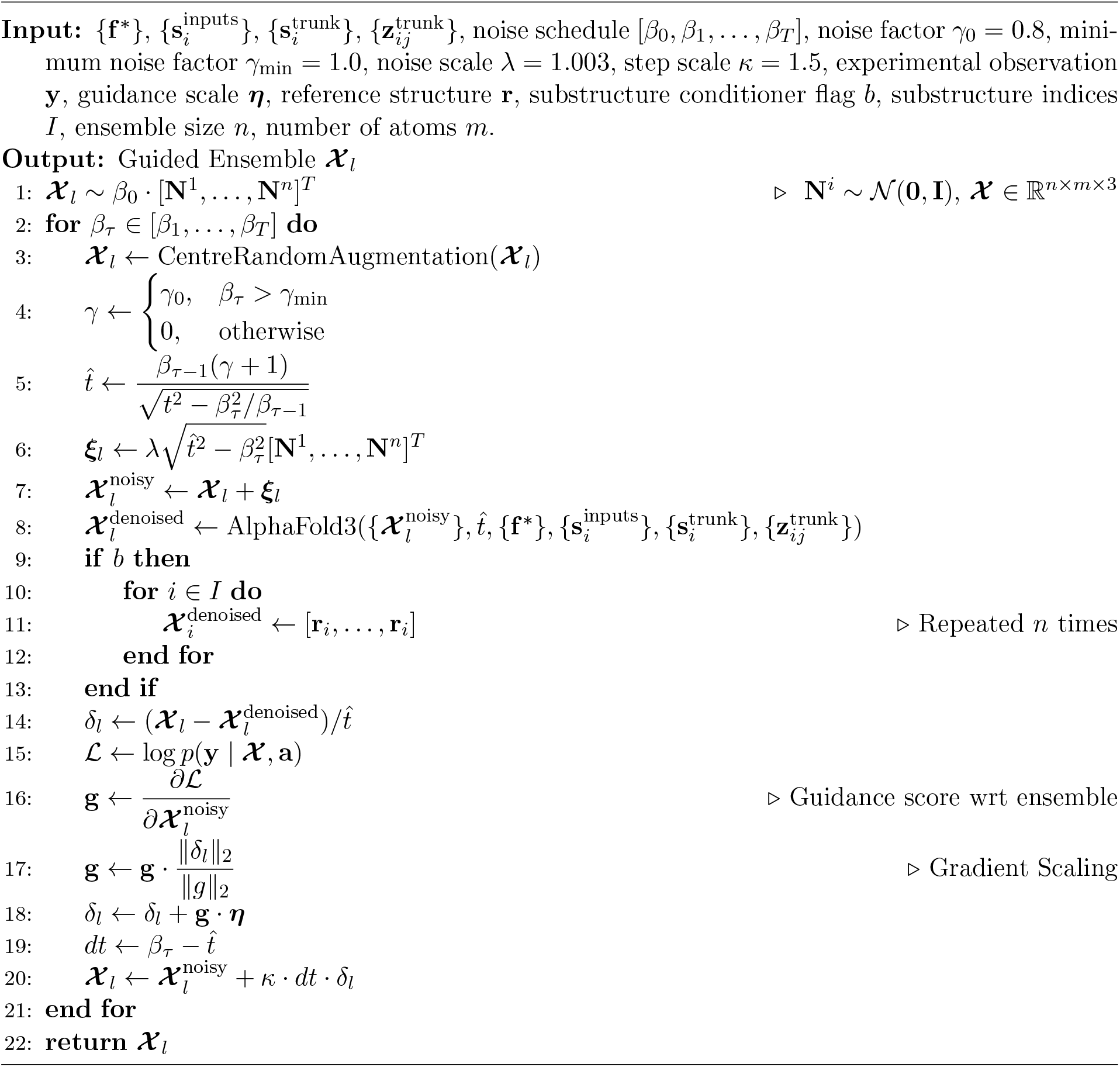

#### Algorithm 2

Likelihood for X-ray crystallography

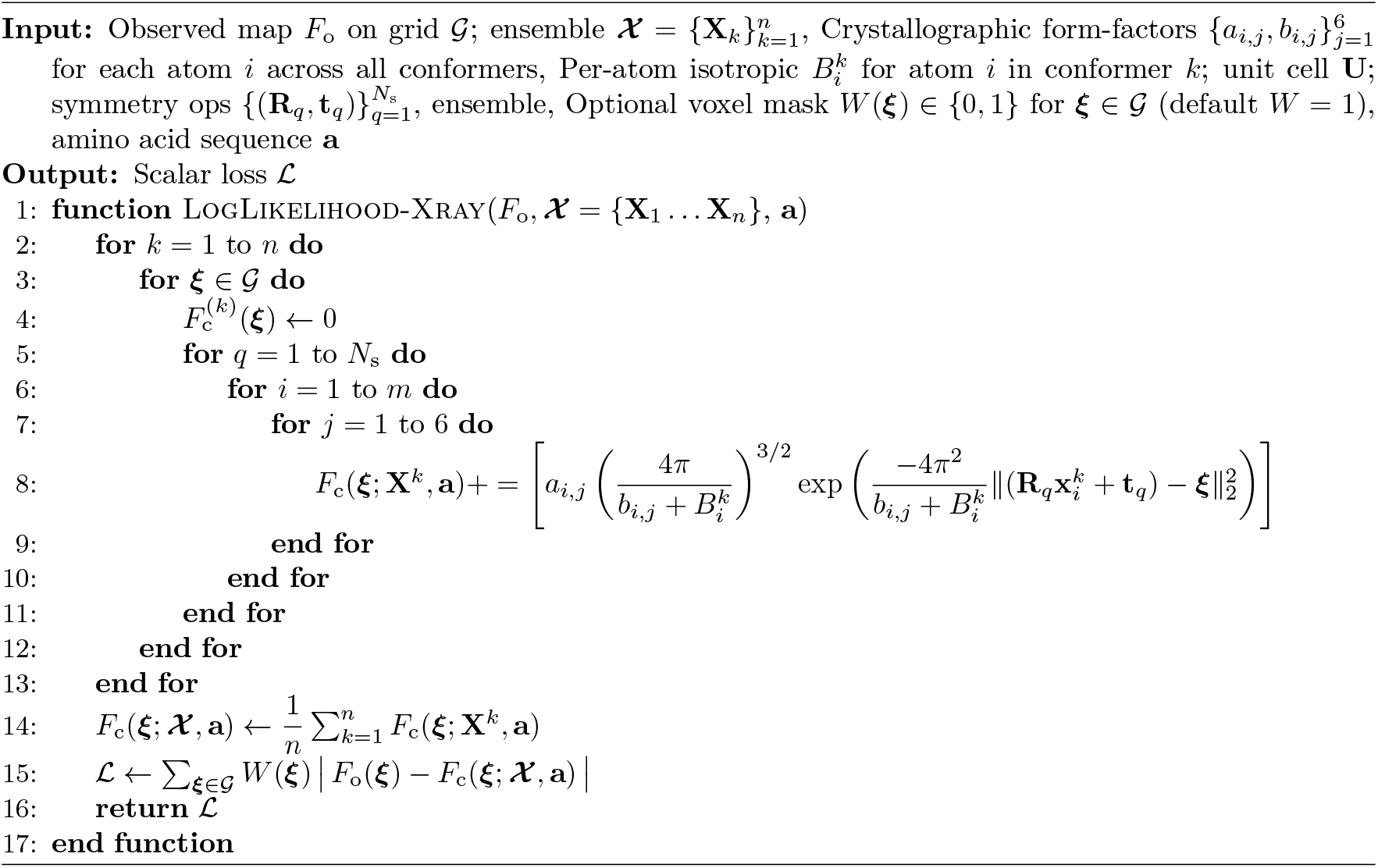

#### Algorithm 3

NOE distance-bound likelihood (ensemble-averaged; ambiguous OR-groups)

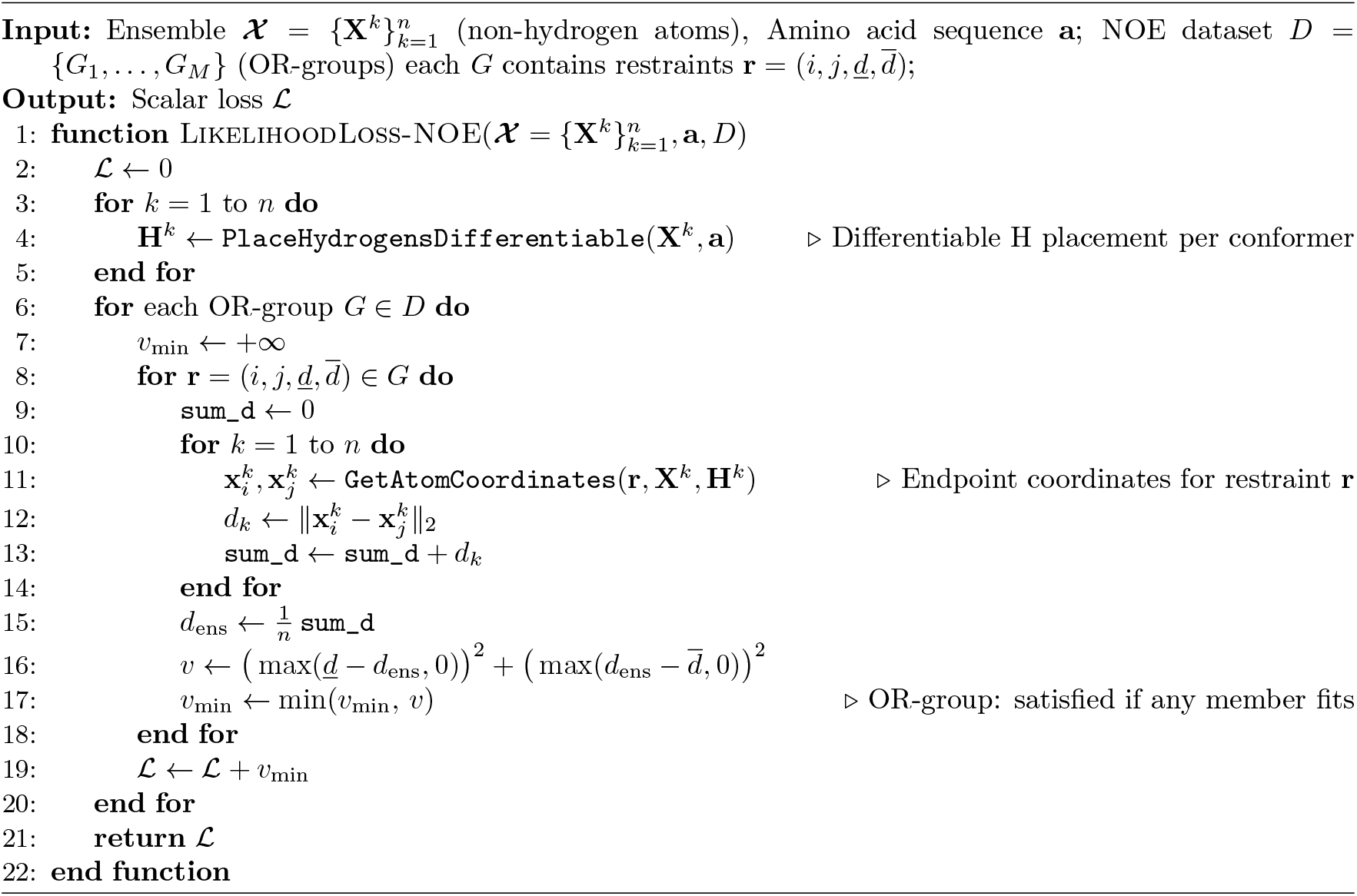

#### Algorithm 4

Order-parameter (*S*^2^) likelihood

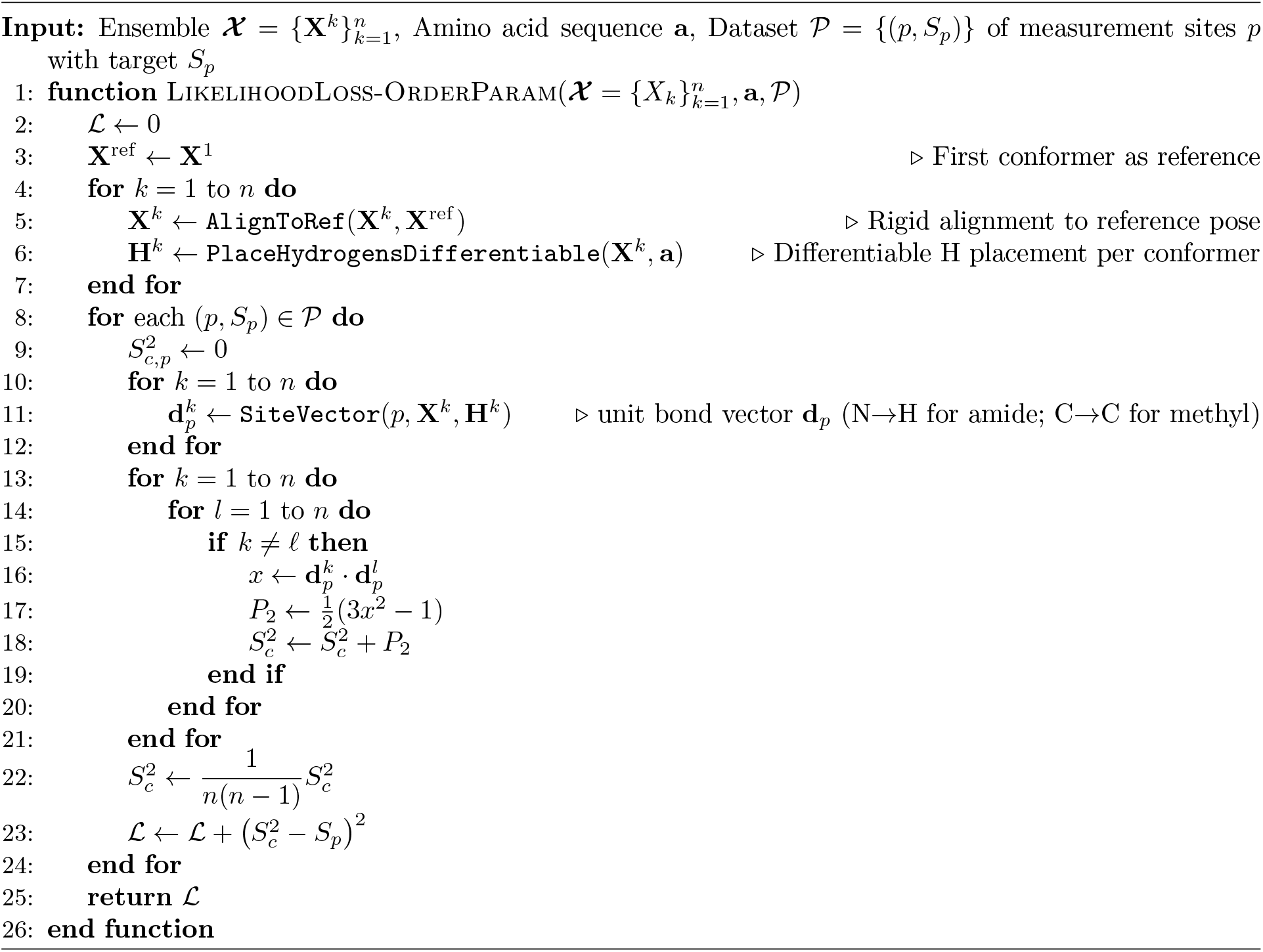

#### Algorithm 5

Selecting samples using matching pursuit [23]

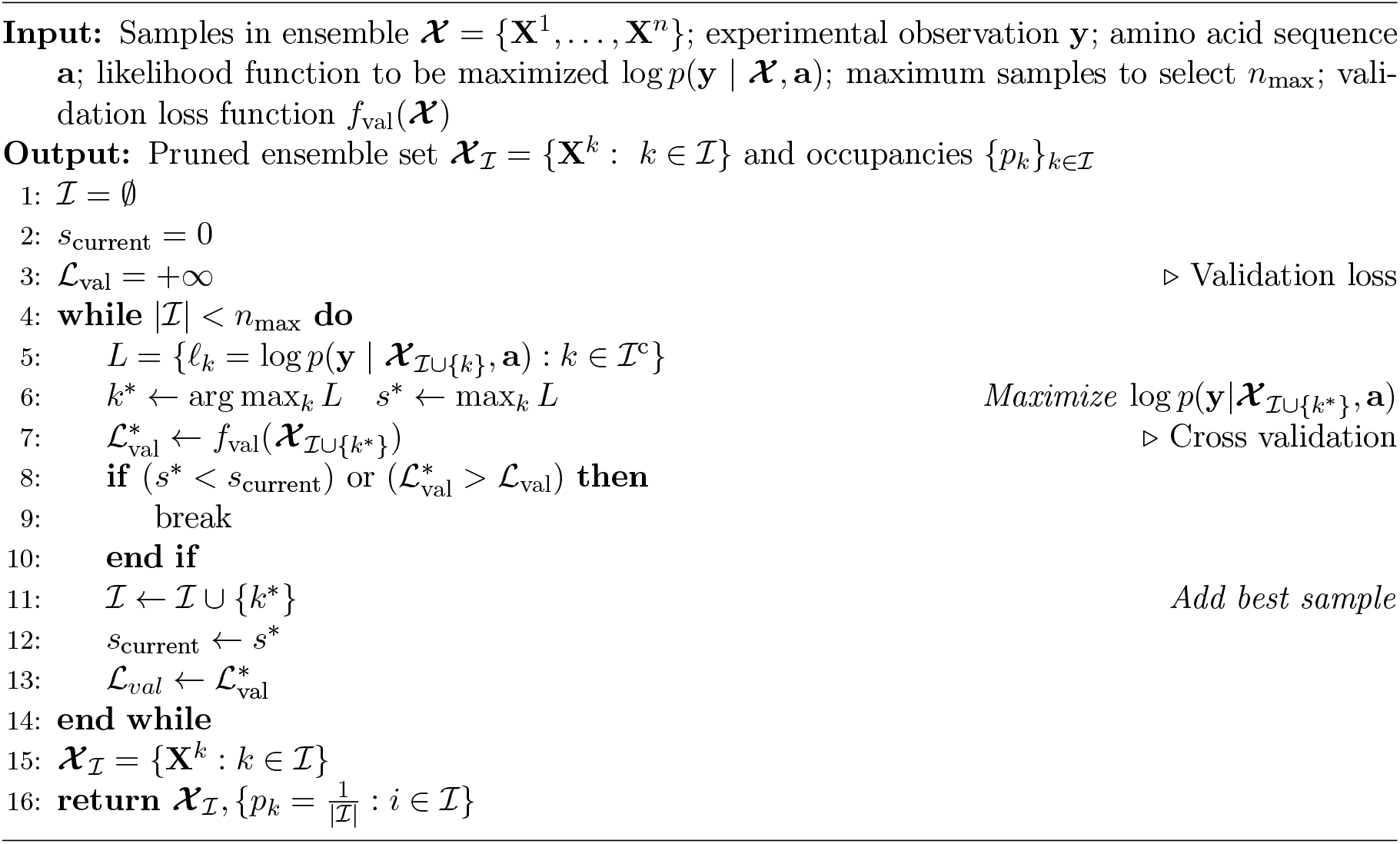

#### Algorithm 6

Occupancy optimization (logit parameterization with entropy sparsity)

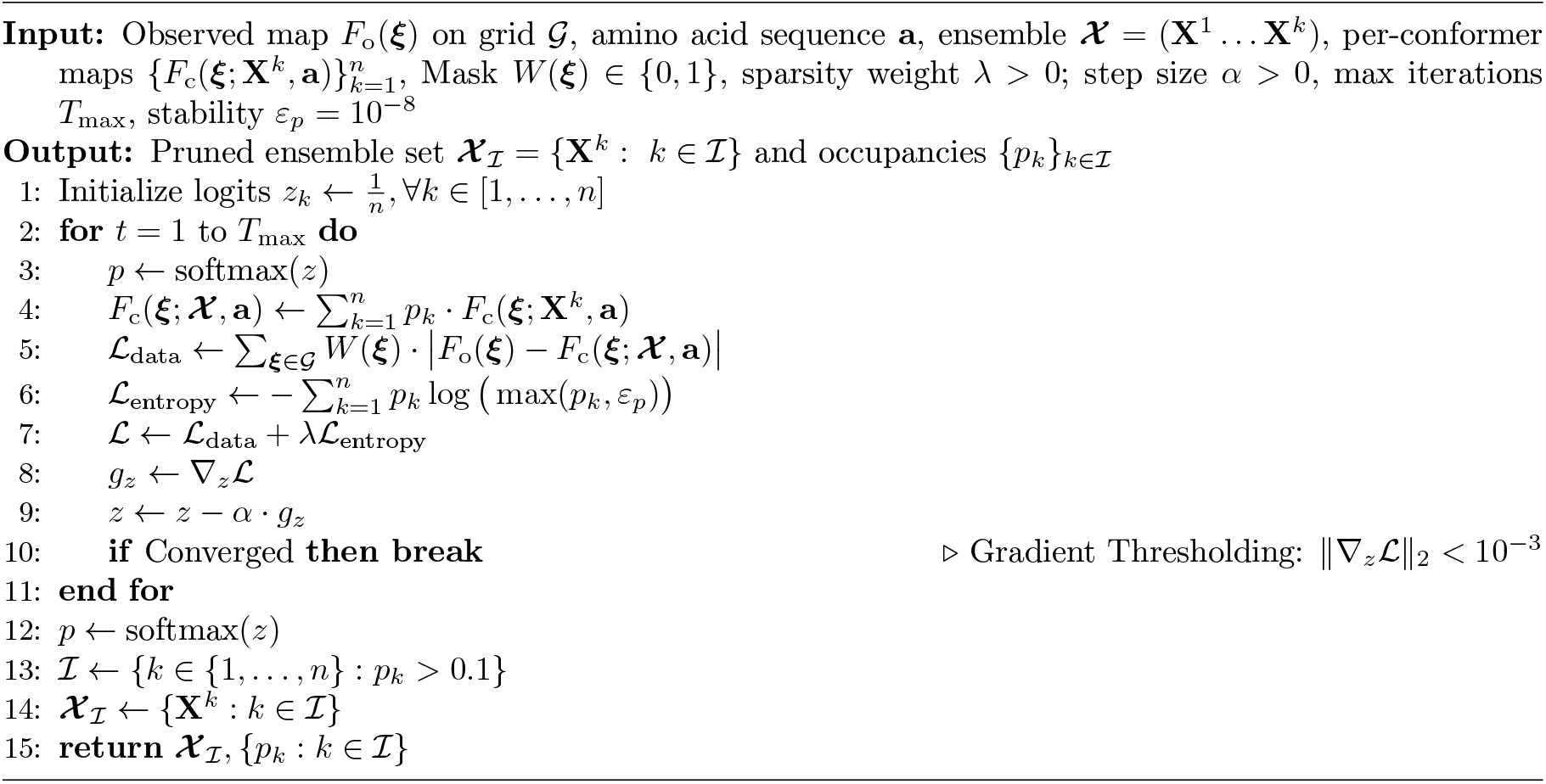

## Footnotes

1 For NOE-guided experiments, we report the best-performing configuration out of two settings.

## Notes

https://doi.org/10.7910/DVN/PLYUHN

https://doi.org/10.5281/zenodo.17307005

### Competing Interest Statement

The authors have declared no competing interest.

### Summary of Updates

Merged SI with the paper and added more tables, plots, and methodological details.

